# Circuit-Specific Early Impairment of Proprioceptive Sensory Neurons in the SOD1^G93A^ Mouse Model for ALS

**DOI:** 10.1101/669788

**Authors:** Soju Seki, Toru Yamamoto, Kiara Quinn, Igor Spigelman, Antonios Pantazis, Riccardo Olcese, Martina Wiedau-Pazos, Scott H. Chandler, Sharmila Venugopal

## Abstract

Amyotrophic Lateral Sclerosis (ALS) is a neurodegenerative disease in which motor neurons degenerate resulting in muscle atrophy, paralysis and fatality. Studies using mouse models of ALS indicate a protracted period of disease development with progressive motor neuron pathology, evident as early as embryonic and postnatal stages. Key missing information includes concomitant alterations in the sensorimotor circuit essential for normal development and function of the neuromuscular system. Leveraging unique brainstem circuitry, we show *in vitro* evidence for reflex circuit-specific postnatal abnormalities in the jaw proprioceptive sensory neurons in the well-studied SOD1^G93A^ mouse. These include impaired and arrhythmic action potential burst discharge associated with a deficit in Nav1.6 Na^+^ channels. However, the mechanoreceptive and nociceptive trigeminal ganglion neurons and the visual sensory retinal ganglion neurons were resistant to excitability changes in age matched SOD1^G93A^ mice. Computational modeling of the observed disruption in sensory patterns predicted asynchronous self-sustained motor neuron discharge suggestive of imminent reflexive defects such as muscle fasciculations in ALS. These results demonstrate a novel reflex circuit-specific proprioceptive sensory abnormality in ALS.

**Significance Statement:** Neurodegenerative diseases have prolonged periods of disease development and progression. Identifying early markers of vulnerability can therefore help devise better diagnostic and treatment strategies. In this study, we examined postnatal abnormalities in the electrical excitability of muscle spindle afferent proprioceptive neurons in the well-studied SOD1^G93A^ mouse model for neurodegenerative motor neuron disease, ALS. Our findings suggest that these proprioceptive sensory neurons are exclusively afflicted early in the disease process relative to sensory neurons of other modalities. Moreover, they presented Nav1.6 Na^+^ channel deficiency which contributed to arrhythmic burst discharge. Such sensory arrhythmia could initiate reflexive defects such as muscle fasciculations in ALS as suggested by our computational model.

## Introduction

Amyotrophic Lateral Sclerosis (ALS) is a neurodegenerative disease in which motor neurons throughout the brain and spinal cord progressively degenerate. In this devastating disease, motor neuron (MN) degeneration leads to muscle paralysis and atrophy, and death ensues 3-5 years following clinical disease onset (Cleveland and Rothstein, 2001; Bruijn et al., 2004). Use of transgenic mouse models of ALS have provided key insights into pre-symptomatic mechanisms of disease development. These include central mechanisms encompassing glutamatergic excitotoxicity at MN synaptic terminals, diminished energy supply resulting from metabolite deficiency, and dysfunctional RNA metabolism, protein homeostasis and aggregation (Bruijn et al., 2004; Taylor et al., 2016). Additional non-cell autonomous contributors such as dysfunctional astrocytes (Yamanaka et al., 2008), microglia (Boillée et al., 2006b), oligodendrocytes (Lee et al., 2012), provide evidence that ALS disease mechanisms are not limited to motor neurons. Peripheral neural dysfunction such as impairment in axonal transport (Williamson and Cleveland, 1999; Puls et al., 2003), and synaptic pruning/failure at the neuromuscular junctions (Pun et al., 2006; Casas et al., 2016) also play a crucial role in initiating muscle paralysis. Importantly, these diverse aspects of early disease development collectively support the idea that additional neuronal pathways such as early sensory input changes are worth exploring as these may provide opportunities for identifying novel early biomarker and therapeutic targets.

Vulnerability to neurodegeneration is not the same for all MNs and within motor pools in ALS (Pun et al., 2006; Kanning et al., 2010). Both in normal aging and in ALS, motor unit (MU) physiology is a crucial determinant of preferential vulnerability, where fast fatigable MUs degenerate first, followed by fast fatigue resistant, while slow MUs are relatively spared (Pun et al., 2006; Hegedus et al., 2007). Secondly, MNs that control eye movements (oculomotor, trochlear and abducens) and sphincter muscles (Onuf’s nucleus), as well as gamma MNs that innervate intrafusal muscle fibers are selectively resistant to death (Iwata and Hirano, 1978; Ferrucci et al., 2010; Lalancette-Hebert et al., 2016). Comparative analyses have highlighted important differences in transcriptional profiles, electrical and synaptic properties, and, neuromuscular biology and innervation patterns between the resistant and vulnerable MNs (Frey et al., 2000; Nimchinsky et al., 2000; Hedlund et al., 2010; Comley et al., 2015; Venugopal et al., 2015; Nijssen et al., 2017). These inherent differences indicate local and long-range defects in sensorimotor circuits of vulnerable MNs in ALS (Durand et al., 2006). For instance, a circuit-specific difference in the ALS-resistant oculomotor neurons involves a lack of Ia muscle spindle afferent proprioceptive inputs (Spencer and Porter, 1988), which are a principal source of glutamatergic excitation during muscle stretch reflexes. A lack of Ia spindle afferent inputs to spinal gamma MNs was suggested as a mechanism of disease resistance in ALS mouse models (Schneider et al., 2012). Consistent with that observation, reducing Ia proprioceptive muscle spindle afferents partly contributed to alpha-MN survival. Therefore, identification and characterization of *circuit-specific* vulnerability to disease progression in ALS (Brownstone and Lancelin, 2018) could help develop effective therapeutic strategies.

To test whether *circuit-specific* dysfunction involves proprioceptive *sensory* neurons, we leveraged the unique architecture of the brainstem trigeminal sensorimotor circuitry involved in jaw control. We examined the proprioceptive Ia afferents in the pontine mesencephalic nucleus (Mes V) in the SOD1^G93A^ mouse model at P11±3 when the jaw motor pools are reported to be dysregulated (Venugopal et al., 2015). Our results show that Mes V neurons present electrical abnormalities and arrhythmic burst discharge patterns, associated with a marked reduction in Nav1.6-type Na^+^ currents. Rescue of these Na^+^ currents restored normal rhythmic burst patterns (also see (Venugopal et al., 2018)). Concomitant examination of trigeminal ganglion neurons and retinal ganglion neurons confirmed exclusive changes only in the proprioceptive Mes V neurons. Using a computational modeling approach we show a functional consequence of sensory abnormality on downstream motor integration predictive of looming reflex dysfunction in ALS.

## Detailed Methods

Transgenic mice expressing high levels of human SOD1^G93A^ (mutant SOD1 or mSOD1) and their wild-type (WT) littermates were used for all the experiments (JAX Strain: 002726 B6SJL-Tg (SOD1*G93A)1Gur/J). All animal protocols were approved by the Institutional Animal Care and Use Committee at UCLA. Experiments were performed at postnatal week 2 (8 - 14 day old mice of either gender) when the rhythmic jaw movements and suckling behavior are fully developed (Turman, 2007). Genotype of mice was determined by standard PCR technique using tails samples (Laragen, Inc, CA). *Experimental preparations and techniques* include: **1)** live brainstem slices to conduct *in vitro* whole-cell current-clamp, voltage-clamp and dynamic-clamp electrophysiology from Mes V sensory neurons, **2)** acutely dissociated trigeminal ganglion neurons to conduct current-clamp experiments, **3)** live whole retinal preparation to conduct current-clamp experiments, **4)** fixed cryosectioned coronal pontine sections for Nav1.6 protein quantification, and, **5)** computational model of Mes V – TMN network to investigate a functional consequence of sensory abnormality on motor discharge.

### I. *In vitro* patch-clamp electrophysiology

#### a. Brainstem slice preparation for Mes V electrophysiology

Brain slices were prepared and used for conducting whole-cell current-, voltage- and dynamic-clamp electrophysiology in the brainstem primary sensory neurons of the trigeminal Mes V nucleus. Pups were anesthetized using isoflurane vapor inhalation, and decapitated. The head was immediately immersed in carboxygenated (95% O_2_-5% CO_2_), ice-cold sucrose cutting solution composed of (in mM): 194 sucrose, 30 NaCl, 4.5 KCl, 1.2 NaH_2_PO_4_, 26 NaHCO_3_, 10 glucose, 1 MgCl_2_. The pontine brainstem was rapidly extracted and adhered to the cutting chamber of a vibratome platform at the rostral end (DSK Microslicer; Ted Pella, Redding, CA); the brainstem was vertically supported by an agar block. The cutting chamber was filled with ice-cold carboxygenated cutting solution. Beginning at the caudal level where the exit of the facial nerve was markedly visible, 3-4 coronal pontine slices, ~250 μm thick were cut and placed in the carboxygenated incubation solution at room temperature, composed of (in mM): 124 NaCl, 4.5 KCl, 1.2 NaH_2_PO_4_, 26 NaHCO_3_, 10 glucose, 2 CaCl_2_, 1 MgCl_2_ (Schurr et al., 1988). The pH of the incubation solution was maintained at 7.28 ± 0.2.

#### b. Trigeminal ganglia extraction and acute dissociation of TGNs for electrophysiology

To evaluate excitability changes in the non-proprioceptive neurons of the trigeminal system of the mSOD1 mice, we performed acute dissociation of the trigeminal ganglia (TG) (Malin et al., 2007; Xu et al., 2010; Yamamoto et al., 2013). Pups were decapitated under isoflurane anesthesia similar to (A) above. The TG were bilaterally removed with the aid of a dissection microscope and transferred into ice-cold (4°C) modified Tyrode’s solution containing (in mM): 130 NaCl, 20 NaHCO_3_, 3 KCl, 4 CaCl_2_, 1 MgCl_2_, 10 HEPES, and 12 glucose, with antibiotic/antimycotic solution (0.5%; Fisher Scientific Company LLC, Hanover park, IL). The ganglia were then minced and incubated in collagenase (1 mg/ml, type I; Fisher) for 40 minutes and then in collagenase with trypsin/EDTA (0.2%; Fisher) for another 40 minutes at 37°C. The TG cells were then washed twice with the modified Tyrode’s solution and triturated gently using fire-polished Pasteur glass pipettes. Finally, the cell suspension was mixed with bovine serum albumin (15%; Fisher) and centrifuged at 900 rpm for 10 min to remove myelin and debris. The pellet was resuspended with Neurobasal A (Fisher) containing B27 (2%; Fisher), L-glutamine (0.2%; Fisher), and antibiotic/antimycotic solution (0.1%), and cells were plated onto glass cover slips coated with Poly-D-lysine/Laminin (Fisher). The cells were then incubated at 37°C in a humidified 5% CO_2_ chamber, and whole-cell patch-clamp electrophysiology was conducted approximately 24 hours after plating (Chen et al., 2008; Marchenkova et al., 2016).

#### c. Retina extraction and preparation for electrophysiology

To ascertain whether visual sensory neurons are susceptible to early changes in excitability in ALS mice, we extracted whole retinas from mSOD1 and WT mice for patch-clamp electrophysiology. For retinal extraction, pups were deeply anesthetized using Isoflurane inhalation and the eyes were denucleated and placed in an ice-cold solution containing (in mM): in room temperature-oxygenated solution containing (in mM): 124 NaCl, 4.5 KCl, 1.2 NaH_2_PO_4_, 26 NaHCO_3_, 10 glucose, 2 CaCl_2_, 1 MgCl_2_. The pH of the incubation solution was maintained at 7.28 ± 0.2, with osmolarity adjusted to 300 ± 5 mOsm. Each retina was exposed by a single cut along the ora serrata. An additional cut was made along the optic disk to dissect the eyecup into two halves (Wang et al., 1997; Qu and Myhr, 2011). The retina was isolated from pigment epithelium and was stored for 1 hour in oxygenated incubation solution at room temperature prior to whole-cell patch-clamp recording.

#### d. Whole-cell current-clamp recording

Whole-cell current-clamp recording was performed in three sets of sensory neurons: the trigeminal proprioceptive Mes V neurons, trigeminal ganglion mechanoreceptive/nociceptive neurons, and the retinal ganglion neurons. Axopatch-1D patch-clamp amplifier (Axon Instruments, Foster City, CA) and pCLAMP acquisition software (version 9.2, Axon Instruments) was used. Silver chloride electrode (Ag/AgCl wire) and 3 mM KCl agar-bridge were used for suitable grounding and electrical isolation. Patch electrodes were fabricated from borosilicate glass capillary tubing (1.5 mm OD, 0.86 mm ID) using a Model P-97 puller (Sutter Instrument, Navato, CA). Tip resistance was 3–5 MΩ when filled with a pipette internal solution containing (in mM): 135 K-gluconate, 5 KCl, 0.5 CaCl_2_, 5 HEPES (base), 5 EGTA, 2 Mg-ATP, and 0.3 Na-ATP (Gupta et al., 2012), pH = 7.25±0.2, osmolarity adjusted to 290 ± 5 mOsm. The external recording solution consisted of (in mM) 140 NaCl, 4 KCl, 2 CaCl_2_, 10 HEPES (base), 2 MgCl_2_, and 10 glucose.

##### i) Mes V neuron identification

The proprioceptive Mes V nucleus was identified bilaterally in the coronal slices as a dorsally located ellipsoid region, ventrolateral to the aqueduct in the caudal pontine slices at the level of VII nerve exit, and in the lateral periphery of the periaqueductal gray in further rostral pontine slices (Wu et al., 2001; Enomoto et al., 2006; Seki et al., 2017). The Mes V neurons were easily distinguished on the basis of their large, ellipsoid soma with ~30-50 *μ*m diameter (Henderson et al., 1982; Del Negro and Chandler, 1997; Enomoto et al., 2006). In a subset of experiments, sensory Mes V neurons were identified using anterograde labeling by injection of 1% Alexa Fluor 568 (Thermo Fisher Scientific) in the jaw closer muscles (20μL on each side) (see **Fig. 1**).

**Figure 1.**
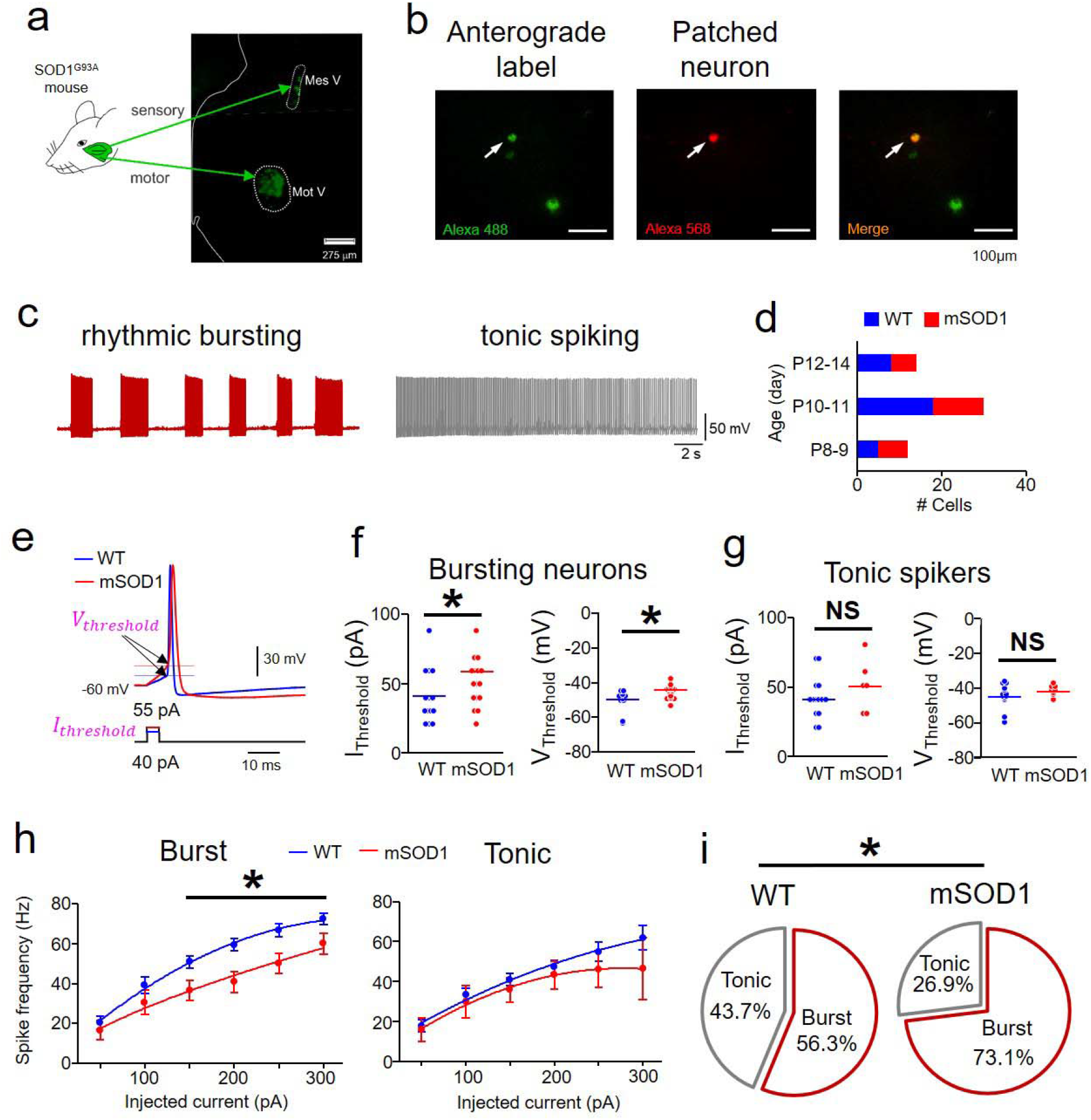
Impaired excitability in intrinsically bursting mSOD1 Mes V neurons. **a.** Schematic shows the Ia proprioceptive sensory and motor nuclei of the trigeminal jaw control circuit identified using antero/retrograde tracer injection in jaw closer muscles in the SOD1^G93A^ mouse (or mSOD1); Left half of a coronal pontine section is shown. Mes V: trigeminal mesencephalic sensory nucleus, Mot V: trigeminal motor nucleus. Both nuclei are demarcated by white dotted enclosures. **b.** Left image shows anterogradely labeled Mes V neurons (green); middle image shows a dye-filled (red) Mes V neuron during patch-clamp recording; right panel shows a merged image highlighting the double-labeled neuron. **c.** Representative examples of discharge patterns in two classes of Mes V neurons; *Left:* a rhythmic bursting neuron (*maroon*). *Right:* a tonic spiking neuron (*right*); These discharge patterns are induced by a 200 pA depolarizing step current for 20 sec duration. **d.** Bar charts summarize the age distribution of the mice including 24 WT mice (*n = 32*) and 21 mSOD1 mice (*n = 26*), where *n* is the number of cells for all the data presented in **Fig. 1**. **e.** A single near-threshold spike in a WT (blue) and a mSOD1 (red) Mes V neuron generated in response to a brief (5 ms) step depolarization; The corresponding injected current threshold (*I_threshold_*) (lower traces) and membrane voltage threshold (*V_threshold_*) for spike generation (upper traces with horizontal lines and arrows) are highlighted (magenta) (see **Methods Section IVb**). **f, g.** Dot plots show *I_t_hreshoid (left panel*), and *V_threshold_* (*right panel*) in bursting (**f**) and tonic (**g**) Mes V neurons; Blue and red horizontal lines indicate average group values. For bursting neurons (**f**), asterisk above black horizontal line indicates *p* < 0.05 for *I_threshold_* and *V_threshold_* in **f**; For tonic spiking neurons (**g**), NS above black horizontal line indicates no statistically significant group differences between WT and mSOD1 cells. **h.** Spike frequency – Injected current responses in bursting (*left*) and tonic spiking (*right*) Mes V neurons: WT (blue) and mSOD1 (red); asterisk above horizontal line indicates *p* < 0.05 between WT and mSOD1 cells for injected current ≥ 150 *pA*. **i.** Pie charts show proportional distribution of bursting and tonic spiking neurons in WT and mSOD1 mice; asterisk indicates *p* < 0.05 in all the panels.

##### ii) Trigeminal ganglion neuron identification

The dissociated TGNs were plated on a coverslip and cells were identified for whole-cell patch-clamp recording using differential interference contrast microscopy. Medium to large isolated cell bodies with diameter > 10μm were selected. During initial experiments, the dissociated cells were labeled using NeuN stain (Neurotrace) for morphology identification and yield estimation. The pipette internal solution was filled with 1% Texas Red (Life Technologies) to label the recorded cells in a subset of experiments (see **Fig. 7**).

##### iii) Retinal ganglion neuron identification

A single retinal quarter was placed in the patch-clamp recording chamber with the ganglion cell layer on the top. Using a recording glass pipette tip, membrane of the surface layer was penetrated and removed by moving the pipette back and forth to expose the retinal ganglion cell layer. The internal limiting membrane encasing the ganglion cell layer was carefully dissected using an empty patch pipette and access was gained to a selected cell for patch-clamp recording (Murphy and Rieke, 2006; Margolis and Detwiler, 2007). Larger cells in the ganglion cell layer (~20 μm diameter) were targeted in the flat-mount retina and recordings were performed in the dark. We did not test cellular response to light versus dark and we performed all our recording in normal ACSF without any synaptic blockers.

#### e. Whole-cell voltage-clamp recording

Voltage-clamp experiments were performed in Mes V neurons to evaluate changes in voltage-gated Na^+^ currents in the mSOD1 mice, compared to their WT littermates as controls. Recordings were performed using a pipette internal solution composed of (in mM): 130 CsF, 9 NaCl, 10 HEPES, 10 EGTA, 1 MgCl_2_, 3 K_2_-ATP, and 1 Na-GTP. The external recording solution consisted of (in mM): 131 NaCl, 10 HEPES, 3 KCl, 10 glucose, 2 CaCl_2_, 2 MgCl_2_, 10 tetraethylammonium (TEA)-Cl, 10 CsCl_2_, 1 4-aminopyridine (4-AP), and 0.3 CdCl_2_. Use of TEA, 4-AP and CdCl_2_ allowed isolation of voltage-gated Na^+^ currents by pharmacologically blocking the voltage-gated K^+^ and Ca^2+^ currents (Del Negro and Chandler, 1997; Enomoto et al., 2006). For acceptable recordings, each cell’s uncompensated series resistance (R_s_) was monitored throughout and only recordings with R_s_ < 10% of the input resistance (R_inp_) were included. Voltage-clamp protocols were used to activate the Nav1.6-type persistent and resurgent voltage-gated Na^+^ currents, both of which are critical for Mes V neuron excitability (Wu et al., 2005; Enomoto et al., 2007). To activate the resurgent Na^+^ current, the voltage-clamp protocol consisted of a brief 3 ms step to +30 mV from an initial holding potential of −90 mV, followed by 100 ms steps from 0 to −90 mV in steps of 10 mV, and then a return to the holding potential of −90 mV (Raman and Bean, 2001; Enomoto et al., 2006). The slower low-voltage-activated persistent Na^+^ current protocol included a slow voltage ramp from −90 mV to +30 mV for 1 second, followed by a step to −40 mV (Do and Bean, 2003; Wu et al., 2005). Following these protocols, these Na^+^ currents were abolished using 0.5μM TTX and leak-subtracted Na^+^ current magnitude was quantified (**Fig. 3**).

#### f. Real-time dynamic-clamp electrophysiology

Real-time closed-loop dynamic-clamp electrophysiology was adopted for *in silico* knock-in of Nav1.6-type Na^+^ currents in Mes V neurons. Briefly, the Linux-based Real-Time eXperimental Interface (RTXI v1.3) was used to implement dynamic-clamp, running on a modified Linux kernel extended with the Real-Time Applications Interface, which allows high-frequency, periodic, real-time calculations (Lin et al., 2010). The RTXI computer interfaced with the electrophysiological amplifier (Axon Instruments Axopatch 200A, in current-clamp mode) and the data acquisition PC, via a National Instruments PCIe-6251 board. Computation frequency was 20 kHz. Brainstem slices were perfused with oxygenated recording solution (~2ml/min) at room temperature (~22 – 24^0^C) while secured in a glass bottom recording chamber mounted on an inverted microscope with differential interface contrast optics (Zeiss Axiovert 10). Current clamp (and dynamic-clamp) data were acquired and analyzed using custom-made software (G-Patch, Analysis) with sampling frequency: 10 kHz; cut-off filter frequency: 2 kHz. The conductance-based Nav1.6-type Na^+^ current models were developed in our lab as detailed in (Venugopal et al., 2018).

#### g. Data acceptance criteria, analysis and statistics

Patch-clamp recording was performed in whole-cell configuration following rupturing of a giga-ohm seal. Cells with uncompensated series resistance > 10 MΩ and resting potential more positive than −50 mV were discarded. Further data acceptance criteria included input resistance ≥ 100 MΩ (measured as the slope, voltage/current at the end of 100 ms current pulses near rest V_rest_ ± 10 mV) and action potential height ≥ 80 mV (measured from spike threshold to peak). Spike characteristics including height, half-width, after-hyperpolarization as well as input threshold (I_threshold_) were determined using response to a 10 ms current pulse. Average spike frequencies used for frequency-current relationships were determined for increasing steps of 1 second current pulses, typically up to 3X I_threshold_. In our Mes V discharge patterns, we distinguished rhythmic bursting activity from tonic spiking by constructing the inter-event-interval (IEI) histogram with bin width of 1 to 5 ms. If the IEI histogram had a clear single peak at around 20 ms, the cells were classified as tonic cells, whereas cells with additional peak(s) at larger IEI values ≥ 100 ms corresponding to the inter-burst intervals or IBIs were grouped as bursting neurons. In the bursting Mes V neurons, we used 40 ms as the cut-off for IEI to further separate the inter-spike intervals (ISIs) from IBIs; all tonically spiking neurons had visibly regular IEIs < 50 ms with a sharp peak in the IEI histograms between 10 – 20 ms. These analyses were performed using Clampfit 10.5; Statistical analyses were performed using Microsoft Excel and Systat 13. Groups were compared using two-sample two-tailed Student’s t-test assuming equal variance, two-way repeated-measures ANOVA to analyze group (WT versus mSOD1) and treatment effects (e.g., different levels of current application) and *χ*^2^ proportionality test for comparison of distribution of cell types in the two animal groups (WT versus mSOD1). A student t-test was used as a posthoc test for 2-way ANOVA. Autocorrelation was used as a measure of rhythmicity of membrane voltage responses in Clampfit 10.5. A p-value < 0.05 was considered statistically significant.

### II. Nav1.6 Fluorescent Immunohistochemistry

#### i) Tissue preparation

Pups were deeply anesthetized using intraperitoneal injection of sodium pentobarbital (2.2 μL/g b.w.) Using a 25-gauge needle connected to a pressure-controlled perfusion system (~200 mmHg) an adequate volume of freshly prepared, chilled 4% Paraformaldehyde (fixative) was transcardially perfused until clear drainage from the right atrium was noted. Subsequently, the mouse was decapitated, and the brain was extracted and incubated in 30% sucrose solution at 4°C for 48 hours. The pontine brainstem block was cut and placed caudal side down within an RNase-free mold and embedded in optimal cutting temperature compound (Tissue TEK-OCT compound, Fisher Scientific), flash frozen using dry ice and stored at −80°C.

#### ii) Cryo-sectioning and fluorescent immunohistochemistry

Pontine brainstem was sectioned at 20 μm thickness beginning approximately at the level of the exiting VII nerve and 6-8 sections per animal were mounted on glass slides (Fisher), ensuring inclusion of the rostro-caudal extent of the Mes V area. Fluorescent immunohistochemistry was immediately performed on slides by marking contours around each section using a MINI PAP PEN (Life Technologies). Every experiment was simultaneously run on WT and mSOD1 pair(s) to ensure similar tissue treatment for subsequent fluorescence quantification. Following three 10-minute washes with 1x PBS, sections were incubated in a blocking buffer (10% normal donkey serum and 0.3% Tween-20 in 1x PBS) for 1 hour at room temperature. Red fluorescence-conjugated Nav1.6 primary antibody derived from rabbit (Alomone Lab) was added (1:200 dilution) and incubated for 24 hours at 4^0^ C. Appropriate dilution was determined based on initial dilution series experiments (1:100, 1:200 and 1:500) with suitable negative controls. Following primary incubation, further wash steps (5 minutes, 3 times in 1x PBS) and green-fluorescence conjugated NeuN stain (Neurotrace) was added (1:200 dilution) for 30 minutes to enable identification of neuronal cell bodies. Sections were rinsed for 5-10 mins with 1x PBS and cover slipped for imaging.

#### iii) Imaging and quantification of Nav1.6 intensities

Imaging was performed using a fully automated epifluorescence microscope (Keyence Bz 9000e). First, the Mes V nuclei were bilaterally identified with NeuN stain at low magnification (2x), ventrolateral to the periaqueductal grey. The identified cells within the nucleus were then imaged at 60x magnification (1/15 s exposure time for NeuN and 2 s exposure time for Nav1.6 with standard excitation settings) and z-stacks were collected at 0.5 *μm* depth resolution with green (NeuN) and red (Nav1.6) filters. The z-projected images were then overlaid to generate a merged image to localize Nav1.6 label on Mes V neurons. Quantification of fluorescent intensities in Mes V neurons was performed using opensource Image J software (Schindelin et al., 2012). First, 12 ± 6 Mes V neurons with clearly visible NeuN stained cell bodies and nuclei were manually traced as regions of interest (ROIs) in each image. These ROIs were then overlaid on the z-projected Nav1.6 image (see **Fig. 4**). The mean ROI intensity and area were quantified for each cell/ROI using the Image J menu-driven functions.

### III. Computational model of simplified sensorimotor network

Our simplified sensorimotor network consisted of conductance-based models for sensory Mes V neuron and trigeminal motor neuron (TMN). All model conductances followed the well-known Hodgkin-Huxley formalism (Hodgkin and Huxley, 1952). The model was implemented using XPPAUT (Ermentrout, 2001) script and will be made available up on request. The model consists of an excitatory network input from the Mes V to the TMN and the model assumptions are detailed below.

The Mes V neuron model consisted of our recently updated version consisting of both persistent and resurgent Na^+^ currents that were reduced in the mSOD1 mouse in this study (Venugopal et al., 2018). Briefly, the model incorporates a minimal set of ionic conductances essential for producing rhythmic bursting and for maintaining cellular excitability in these neurons (Wu et al., 2005). These include a potassium leak current, *I_leak_*, a 4-AP sensitive delayed-rectifier type potassium current *I_K_*, and a Nav1.6-type sodium current, *I_Na_* with three separable components: fast, persistent and resurgent currents. Each of these model currents are based on our previous experimental work (Chandler et al., 1984; Del Negro and Chandler, 1997; Wu et al., 2001; Wu et al., 2005; Enomoto et al., 2006; Venugopal et al., 2018).

The TMN model is based on well-established two-compartment MN models which include dendritic Ca^2+^ currents (e.g., (Booth et al., 1997; Venugopal et al., 2011a)). This is based on known physiology of MNs which richly express non-inactivating dendritic L-type Ca^2+^ currents, also known as the persistent Ca^2+^ current (CaP) in both the brainstem and spinal cord (Lee and Heckman, 1998; Carlin et al., 2000; Theiss et al., 2007). We chose such a model to reproduce dendritic plateau potentials, membrane bistability (Booth et al., 1997) and also since the CaP plays a crucial role in synaptic amplification (Hultborn et al., 2003) and motor control (Lee et al., 2003; Heckman et al., 2005). The somatic compartment consisted of the action potential generating fast sodium, delayed-rectifier type potassium, high-voltage activated N-type Ca^2+^ and N-type Ca^2+^ activated potassium conductances (Hsiao et al., 1998; Hsiao et al., 2005). The plateau generating L-type persistent Ca^2+^ or CaP, CaP-activated potassium and a persistent sodium conductance were confined to the dendritic compartment. Both compartments also consisted of a potassium leak conductance.

A lumped synaptic variable modeled the excitatory synaptic input from the Mes V to TMN model. This assumption was based on known anatomy and physiology of Mes V-TMN connections. For example, nearly 80% of Ia muscle spindle afferents from the Mes V nucleus synapse on to proximal and distal dendrites of TMNs (Yoshida et al., 2017). To incorporate such dendritic excitation, we modeled the TMN with two electrically coupled morphological compartments (soma and lumped dendrite) (Booth et al., 1997; Venugopal et al., 2011a; Venugopal et al., 2012) and confined synaptic input to the dendritic compartment. Secondly, we tuned the strength of the model excitatory conductance to generate sensory-coupled spikes in the TMNs since Mes V neurons provide strong monosynaptic glutamatergic excitation to TMNs (Chandler, 1989; Del Negro and Chandler, 1998; Turman et al., 1999; Turman et al., 2000). For simplicity, we assumed no other form of synaptic excitation or inhibition. This allowed us to exclusively examine the consequence of irregular sensory patterns as observed in our data, on motor neuron discharge. The lumped synaptic variable represented a classic excitatory synapse modeled as follows (Venugopal et al., 2011b):

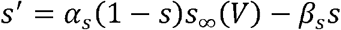

where, *s* is the synaptic variable, *α_s_* = 2 is the rate constant for fast rise of *s, β_s_* = 0.05 is the decay rate constant for relatively slower decay of *s* (Powers et al.; Heckman and Binder, 1991; Trueblood et al., 1996). The steady-state voltage-dependent function, *s*_∞_(*V*) is given by,

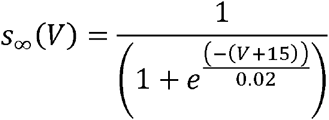

where, *V* is the voltage of the presynaptic neurons (here, Mes V neuron). The excitatory postsynaptic current was included in the dendritic compartment of the TMN model and is given by the following conductance-based equation:

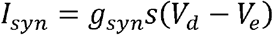

where, *V_d_* is the TMN dendritic voltage, and, *V_e_* = −20 mV, is the reversal potential for an excitatory synaptic current. The value of maximal synaptic conductance, *g_syn_* was set such that presynaptic Mes V bursts generated postsynaptic TMN firing frequencies in the range observed in SOD1^G93A^ TMNs (Venugopal et al., 2015).

## Results

### Impaired excitability of SOD1^G93A^ proprioceptive Mes V neurons

First, we examined whether the electrical excitability of the jaw muscle proprioceptive neurons in the trigeminal Mes V nucleus show abnormalities in the mSOD1 mice. Using suitable morphological and anatomical selection criteria, we conducted *in vitro* whole-cell patch-clamp recording and evaluated the basic membrane properties, action potential characteristics and responses to increasing step depolarizations. The comparative membrane properties in mSOD1 and WT are summarized in **Tables 1, 2**. In **Fig. 1a**, we illustrate the location of the anterogradely labeled jaw closer muscle spindle afferents in the brainstem Mes V nucleus (green), obtained from injection of dye into the jaw closer muscles. Concomitantly, the jaw closer motor pools are also retrogradely labeled specifically in the dorsolateral trigeminal motor nucleus (Mot V) as shown in **Fig. 1a**, ventrolateral to Mes V. In **Fig. 1b**, we present an example of an anterogradely labeled Mes V sensory neuron, which is dye-filled during whole-cell patch-clamp recording. **Figure 1d** shows the age distribution of mice from which our dataset was obtained. In **Fig. 1c**, we show representative examples of the two most commonly observed patterns of action potential discharges in our dataset from both WT and mSOD1 mice (Brocard et al., 2006; Enomoto et al., 2006). These included: **1)** rhythmic burst discharge consisting of repetitive sequences of high frequency spikes (~80 Hz), followed by periods of quiescence on the order of 100 – 1000 ms (Wu et al., 2001; Brocard et al., 2006), and, **2)** a tonic spiking pattern with short inter-spike intervals, on the order of 10 ms. Our empirical criteria for classifying cells as bursters was based on the distributions of spike intervals (see **Methods Section I.g**). In the mSOD1 bursting Mes V neurons, we noted marked increase in the current and voltage thresholds for spike generation (see **Fig. 1e**, and **Fig. 1f**) compared to the WT. The mean ± s.d. for *I_threshold_* were 40.7 ± 5.4 pA and 57.1 ± 7.2 pA, for *V_threshold_* were −50.1 ± 2.1 mV, and −44.6±1.1 mV for WT and mSOD1 respectively. Corresponding *p* values from a two-sample student t-test were 0.04 for *I_threshold_* and 0.008 for *V_threshold_* comparisons. The bursting neurons in the mSOD1 mice also showed diminished spike frequencies in response to increasing current injections (see **Fig. 1h**). A two-way ANOVA statistic showed significant group versus treatment effects (*p* < 0.05). A post-hoc student t-test further confirmed significant reduction in spike frequencies in the mutant for all the current injections ≥ 150 pA (Student’s t test, 150 pA: *p* = 0.0123, 200 pA: *p* = 0.0035, 250 pA: *p* = 0.075, 300 pA: *p* = 0.022). Furthermore, a greater proportion of mSOD1 Mes V neurons presented burst-like activity (**Fig. 1i**). A chi-square proportionality test yielded *p* = 0.0074. Conversely, tonically spiking neurons were less perturbed and did not present any *I_threshold_* or *V_threshold_* changes; however, with higher current injections ≥ 250 pA, these neurons also showed reduced firing frequencies similar to bursting neurons (Student’s t test, 250 pA: *p* = 0.0125, 300 pA: *p* = 0.0361) (see **Fig. 1e**, and **Fig. 1g**). We noted that these neurons also showed an increase in spike width (see summary **Tables 1** and **2**). Moreover, the proportion of tonically firing cells was reduced in the mSOD1 Mes V dataset (**Fig. 1i**). Taken together, Mes V neurons in the SOD1^G93A^ mouse are hypoexcitable with a large propensity towards burst formation in action potential trains.

**TABLE 1.** Membrane properties of Mes V neurons.

|  | Burst |  | Tonic |  |
| --- | --- | --- | --- | --- |
|  | WT (n=18) | mSOD1 (n=19) | WT (n=14) | mSOD1 (n=7) |
| RMP, mV | -56.9 ± 1.5 | -54.8 ± 1.0 | -53.6 ± 1.6 | -52.8 ± 1.4 |
| $R_{in}$, MΩ | 211.5 ± 25.9 | 253.1 ± 19.3 | 264.2 ± 16.9 | 300.1 ± 18.2 |
| $C_m$, pF | 49.2 ± 3.1 | 50.3 ± 2.7 | 42.0 ± 2.1 | 42.2 ± 2.2 |

**TABLE 2.**
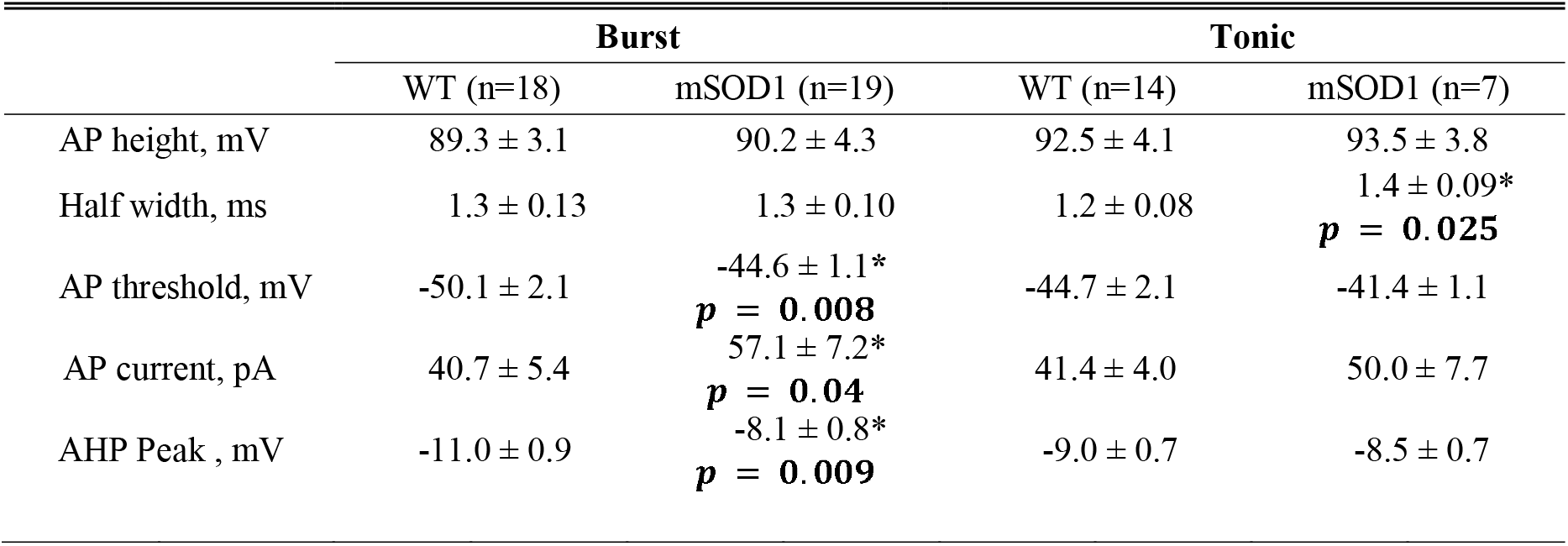
Action potential properties of Mes V neurons (p values are based on student t-test)

|  | Burst |  | Tonic |  |
| --- | --- | --- | --- | --- |
|  | WT (n=18) | mSOD1 (n=19) | WT (n=14) | mSOD1 (n=7) |
| AP height, mV | 89.3 ± 3.1 | 90.2 ± 4.3 | 92.5 ± 4.1 | 93.5 ± 3.8 |
| Half width, ms | 1.3 ± 0.13 | 1.3 ± 0.10 | 1.2 ± 0.08 | 1.4 ± 0.09* |
|  |  |  |  | **p = 0.025** |
| AP threshold, mV | -50.1 ± 2.1 | -44.6 ± 1.1* | -44.7 ± 2.1 | -41.4 ± 1.1 |
|  |  | **p = 0.008** |  |  |
| AP current, pA | 40.7 ± 5.4 | 57.1 ± 7.2* | 41.4 ± 4.0 | 50.0 ± 7.7 |
|  |  | **p = 0.04** |  |  |
| AHP Peak , mV | -11.0 ± 0.9 | -8.1 ± 0.8* | -9.0 ± 0.7 | -8.5 ± 0.7 |
|  |  | **p = 0.009** |  |  |

### Arrhythmic spike and burst patterns of SOD1^G93A^ Mes V neurons

This overall reduction in action potential generation capability and increased burst discharge in the mSOD1 Mes V neurons was accompanied by further abnormalities in bursting properties. In **Fig. 2a**, we illustrate a rhythmic burst pattern in a WT Mes V neuron (*left panel*) compared to irregular burst patterns in a mSOD1 Mes V neuron (*right panel*). The corresponding IEI time series highlight such irregularities in **Fig. 2b** (see legend). Note that in the rhythmic WT burst pattern, the IBIs are well demarcated from the ISIs. However, this distinction is less obvious in the mutant pattern and as noted, the IBI regularity from one burst to another is also uncertain. The ISIs also presented significant irregularities observed for the shorter spike intervals. We used the autocorrelation function of the membrane voltage and compared its 2^nd^ peak between WT and mSOD1 cells using a time lag, *τ* = 100 *ms* to examine regularity of spikes (Rieke et al., 1999). This time window allowed examination of inter-spike irregularities within a typical burst. **Figure 2c** illustrates representative examples of the autocorrelation function for WT (*left*) and mSOD1 (*middle*) Mes V neurons. A two-sample unpaired student t-test with assumed equal variance showed that the 2^nd^ peak of autocorrelation between the two groups was significantly different. A *p* = 0.0029 was noted with mean WT peak ± s.d. as 0.44 ± 0.04, and the mean mSOD1 peak ± s.d. as 0.30 ± 0.03. We also note that the subsequent peaks are almost completely suppressed further highlighting irregularity in spike patterns in the mutant. Additionally, the burst duration and IBI lengths were reduced while the ISIs were increased (see **Fig. 2d**): for WT and mSOD1 respectively, the mean BD ± s.d. were 0.41±0.11 s and 0.06±0.01s, mean IBI ± s.d. were 0.89±0.14 s and 0.29±0.03 s, and mean ISI ± s.d. were 17.2±0.34 ms and 26.4±2.37 ms. Taken together, the proprioceptive Mes V neurons present early impairment in action potential generation and in the ability to sustain regular burst discharge in the SOD1^G93A^ mouse.

**Figure 2.**
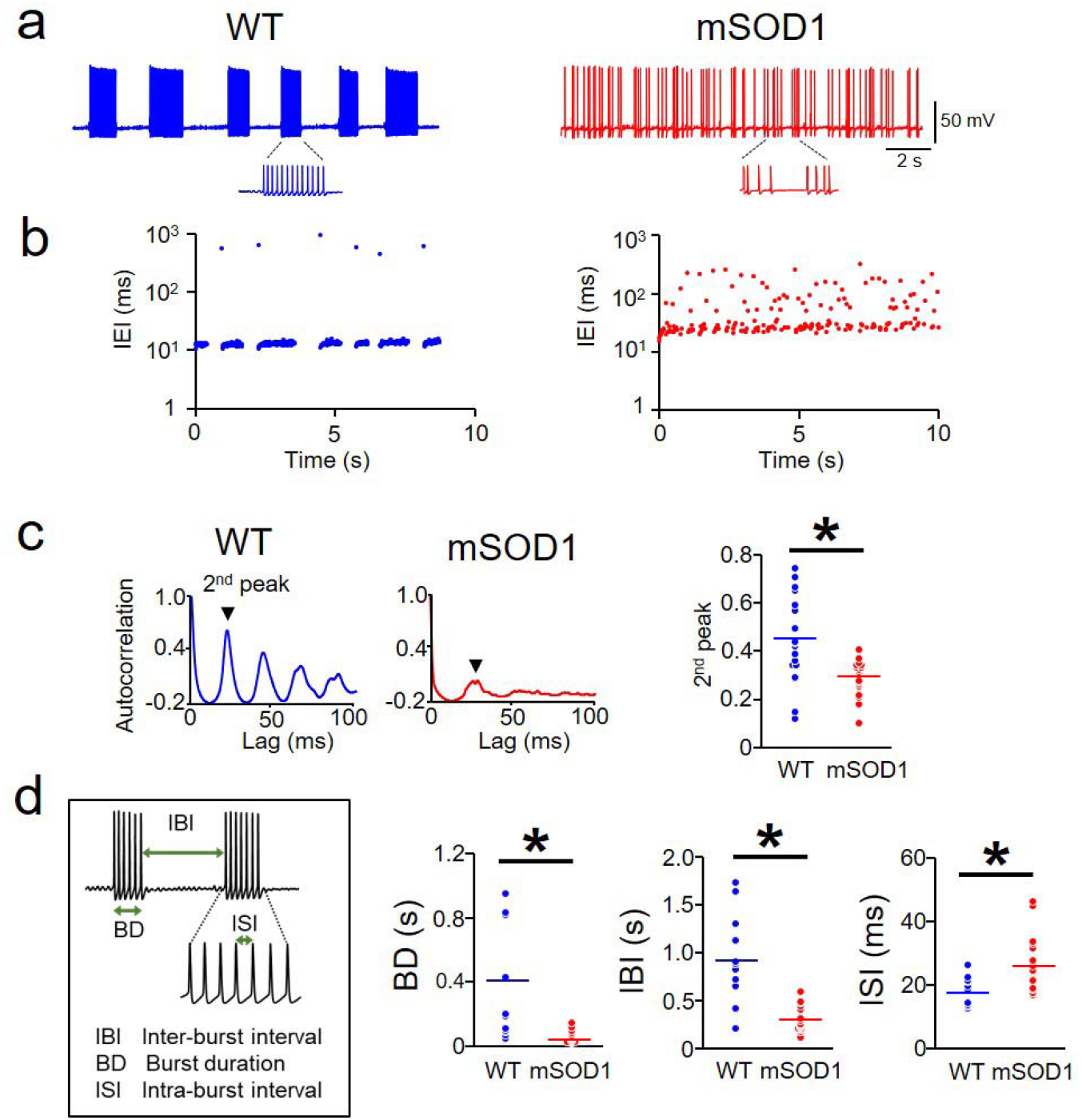
Arrhythmic burst discharge in mSOD1 Mes V neurons. **a.** Representative examples showing rhythmic bursting in a WT (blue) Mes V neuron (*left panel*), and arrhythmic bursting in a mSOD1 (red) Mes V neuron (*right panel*). **b.** Time series plots of IEIs corresponding to the examples shown in (**a**) highlight IEI irregularities (IEIs are shown on log scale). **c.** Autocorrelation function of the membrane voltage in a WT (*left panel*) and mSOD1 Mes V neuron (*middle panel);* 2^nd^ peak is highlighted; Dot plots on the right summarize the 2^nd^ peaks in WT (blue) and mSOD1 (red) bursting neurons. **d.** *Left:* Inset shows the timing properties of bursts. Dot plots show significantly reduced burst duration (BD), inter-burst intervals (IBI) and increased inter-spike intervals (ISI) in the mSOD1 (red) Mes V neurons compared to WT (blue). For data in panels (**c**) and (**d**), *n* = 14 (WT) and *n* = 16 (mSOD1), where, *n* is the number of bursting cells. The asterisk above the black horizontal line indicates a *p* < 0.05 in all the panels.

### Disrupted firing patterns of SOD1^G93A^ proprioceptive Mes V neurons are linked to reduced Nav1.6 Na^+^ currents and channel protein

To identify the ionic basis for the observed impairments in action potential bursts, we tested whether the voltage-gated Na^+^ currents which are essential for spike generation and burst control in these neurons are compromised (Wu et al., 2005; Enomoto et al., 2006; Yang et al., 2009). Our previous report using the Nav1.6 subunit knock-out mouse demonstrated significant reductions in TTX-sensitive Na^+^ currents which paralleled impairment in spike discharge (Enomoto et al., 2007). These Na+ currents including the persistent and resurgent components are important for generation and maintenance of burst discharge in Mes V neurons (Wu et al., 2005; Enomoto et al., 2006; Venugopal et al., 2018). Therefore, we examined whether the observed impairment in both excitability and burst discharge in a large majority of the mSOD1 Mes V neurons could be explained by reductions in the functional expression of these Nav1.6 Na^+^ channel currents. To test this, we used the *in vitro* voltage-clamp approach combined with suitable pharmacology and measured the Nav1.6 channel mediated persistent and resurgent Na^+^ currents in WT and mSOD1 Mes V neurons (see **Methods Section IIb**). As shown in **Figs. 3a – d**, the mSOD1 Mes V neurons showed significantly reduced current density (pA/pF) of voltage-gated persistent and resurgent Na^+^ current components, compared to WT. As shown in **Fig. 3c**, for persistent Na^+^ current, a two-way ANOVA showed group (WT and mSOD1) versus treatment (voltage command steps) effect (*p* = 0.0053) where the mean ± s.d. at −60 mV were - 1.01±0.22 pA/pF and −0.43±0.10 pA/pF, and at −50 mV were 1.15±0.21 pA/pF and −0.66±0.11 pA/pF for WT and mSOD1 cells respectively. Corresponding post hoc student t-test *p* values at - 60 and −50 mV were 0.03 and 0.044 respectively. Similarly, as shown in **Fig. 3d**, for resurgent Na^+^ current, a two-way ANOVA showed group versus treatment effect (*p* = 0.0191) where the mean ± s.d. at −50 mV were −5.13±0.282 pA/pF and −3.23±0.51 pA/pF and at −40 mV these values are −5.69±0.84 pA/pF, and −3.93±0.47 pA/pF for WT and mSOD1 cells respectively. Corresponding post hoc student t-test *p* values at −50 and −40 mV are 0.0387 and 0.0424, respectively. Such reductions in current density, however, were not accompanied by altered voltage-gating properties in the measured Na^+^ currents between WT and mSOD1 Mes V neurons, as noted by the normalized conductance Boltzmann curves (see **Figs. 3e, f**).

**Figure 3.**
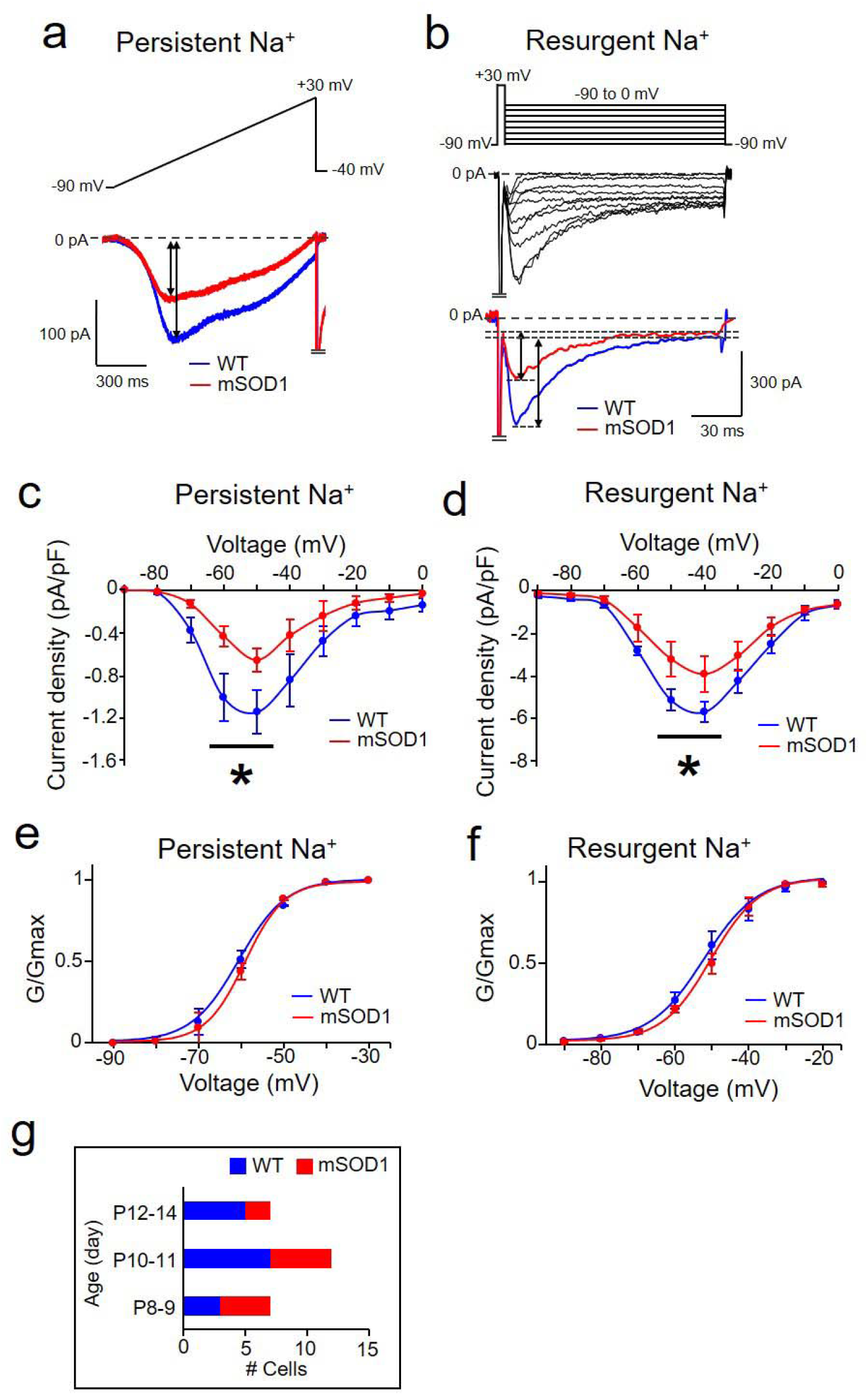
Reduced Nav 1.6-type Na^+^ currents in mSOD1 Mes V neurons. **a, b.** Representative current traces from whole-cell voltage-clamp experiments showing a reduction in TTX-sensitive persistent Na^+^ (**a**) and resurgent Na^+^ (**b**) currents in mSOD1 Mes V neuron (red), compared to WT (blue); top traces show the voltage-clamp protocol and bottom traces show current responses. Voltage-clamp protocol in (**a**) consists of a ramp voltage from −90 to +30 mV at 100 mV/s, and in (**b**), consists of a brief, 5 ms step from −90 to +30 mV, to evoke a transient Na^+^ current, followed by repolarization to voltages between −90 to 0 mV to elicit resurgent Na^+^ current due to open-channel unblock. **c, d.** Current-voltage relationships of the peak persistent (**c**) and resurgent (**d**) Na^+^ current densities in mSOD1 Mes V neurons (red). In (**c**), asterisk above the black horizontal line indicates statistically significant reduction in peak current amplitudes at sub-threshold activation voltages between −60 and −50 mV for persistent Na^+^ peaks in mSOD1 (red) compared to WT (blue). In (**d**), asterisk above the black horizontal line indicates statistically significant reduction in peak current amplitudes at repolarization voltages of −50 and −40 mV for resurgent Na^+^ peaks in mSOD1 (red) compared to WT (blue). **e, f.** Boltzmann curves fit to the normalized peak conductances as a function of activation voltages for persistent (**e**), and resurgent (**f**) Na^+^ currents in WT (blue) and mSOD1 (red) Mes V neurons. **g.** Bar charts summarize the age distribution of the mice including 8 WT mice (*n = 14*) and 6 mSOD1 mice (*n = 11*), where *n* is the number of cells for all the data presented in panels **c – f.**

**Figure 4.**
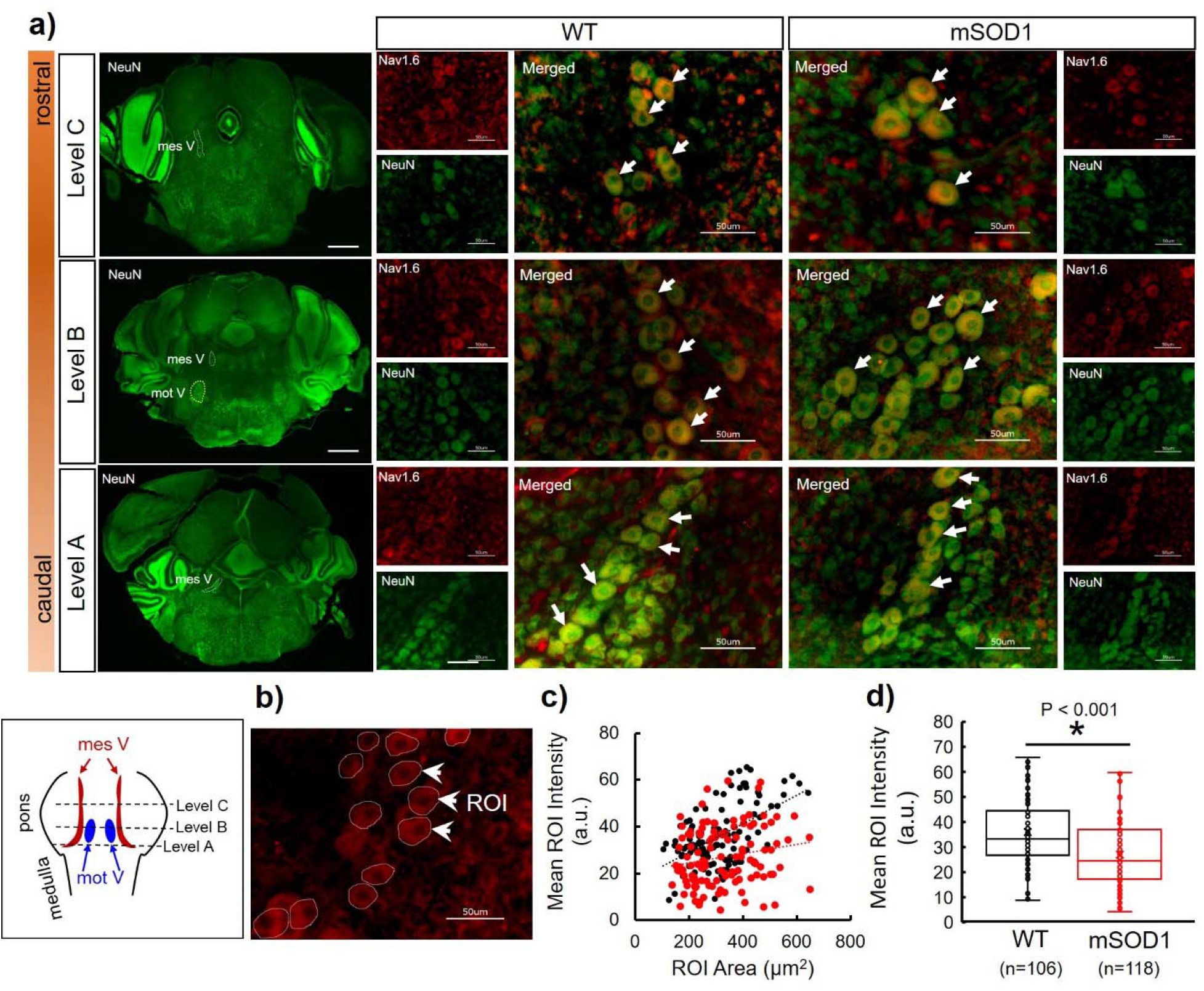
Immunofluorescent quantification of Nav1.6 protein expression in WT and mSOD1 Mes V cells. **a)** Representative images showing three rostro-caudal levels of coronal brainstem sections, stained with NeuN (left column of images); scalebar shows 500 μm. In these images, the Mes V nucleus is highlighted with dashed contours. At Level B, the subjacent trigeminal motor nucleus (mot V) is highlighted. For each level, comparative images consisting of Mes V neurons from WT and mSOD1 mice are shown at 60x magnification; green is NeuN, red is Nav1.6 protein. Merged images are shown enlarged with white arrows highlighting representative Mes V neurons. Left bottom boxed inset illustrates the pontine levels at which sections were collected. **b)** Representative image showing regions of interest (ROIs) drawn around 12 Mes V neurons for immunofluorescence quantification; white arrows highlight three representative ROIs **c)** Scatterplot showing WT (black circles) and mSOD1 (red circles) values of ROI area and mean intensity quantification. Trendlines show a positive linear regression. **d)** Box plots show mean ROI intensity per cell for WT (black) and mSOD1 (red) respectively; *n* values indicate number of cells obtained from 4 WT and 4 mSOD1 mice across 9 and 8 sections respectively. A two-tailed Student t-test was used for statistical comparison.

Next, using fluorescent immunohistochemistry, we tested whether the observed decrease in the Nav1.6 Na^+^ currents in the mSOD1 mouse could result from a down-regulation of Nav1.6 Na^+^ channels on Mes V neuronal membrane (Enomoto et al., 2007). We quantified the Nav1.6 protein expression in Mes V neuron cell bodies colocalized with NeuN stain. The NeuN-stained Mes V neurons were readily detected based on their oval cell bodies and rostro-caudal distribution in the dorsal pontine sections, ventrolateral to the periaqueductal grey (see **Fig 4a** and **inset**). As noted in the figure, the Nav1.6 protein expression was discernable in both WT and mSOD1 Mes V neurons. The three NeuN images (green) shown as level A, B and C illustrate the representative caudal-to-rostral sections used in our analysis. For immunofluorescent quantification, we manually outlined the NeuN-stained Mes V cell bodies (regions-of-interest or ROIs) in which the nuclei were clearly visible under high magnification (60x) (see **Methods**). These ROIs were then overlaid on the Nav1.6 protein-stained image (see **Fig 4b**) and the ROI areas and mean intensities were quantified using the open-source Image J software (Schneider et al., 2012). **Figure 4c** shows a scatterplot of ROI areas and mean intensities for WT (black circles) and mSOD1 (red circles) Mes V neurons to illustrate that there was no sampling bias in our cell samples. Note that there was an overall positive correlation between ROI/cell area and mean intensities in both WT and mSOD1 Mes V neurons (trend lines have positive slopes in **Fig. 4c**). However, such a correlation was weaker in the mutant. Statistical comparison of ROI areas and mean intensities between WT and mSOD1 revealed significantly diminished mean intensities in the mutant (mean ± std: 36.02 ± 13.18 a.u. for WT and 27.01 ± 12.61 a.u. for mSOD1; *p* < 0.001 using two-sample Student t-test) (see **Fig. 4d**; box plots show 1^st^, 2^nd^ and the 3^rd^ quartiles and error bars show 1.5x deviations from the median). However, the WT and mSOD1 ROI areas were not statistically different (not shown) whereas, the mean intensities per unit area (a.u./*μm*^2^) were also diminished in the mutant (*p* < 0.001). These results indicate that the reduced functional expression of Nav1.6 Na^+^ currents in the mSOD1 Mes V neurons are associated with a down-regulation of Nav1.6 protein expression.

### Rescue of Nav1.6 Na^+^ currents restored normal burst patterns and rhythmicity in SOD1^G93A^ proprioceptive Mes V neurons

To test whether rescuing the Nav1.6 Na^+^ current impairment can restore the normal burst discharge in the mSOD1 Mes V neurons, we used an *in vitro* dynamic-clamp approach. This included real-time injection of conductance-based Nav1.6 Na^+^ currents into a mSOD1 Mes V neuron during whole-cell current-clamp recording of burst discharge (see **Fig. 5a**). **Figure 5b** illustrates that addition of such realistic computer-generated Nav1.6 Na^+^ persistent and resurgent currents into an irregularly bursting mSOD1 Mes V neuron converted an irregular burst pattern to a more rhythmic pattern. The red traces in **Fig. 5b** show the membrane voltage of the mSOD1 Mes V neuron during default burst discharge, and the black trace in between shows the modified burst discharge upon addition of the Nav1.6-type Na^+^ currents. To evaluate whether the rescue of Nav1.6-type Na^+^ currents in the mSOD1 neurons also restored the normal burst patterns similar to WT, we quantified the time intervals between spikes (IEIs) and demonstrate that the irregularities in mSOD1 IEIs were abolished (see **Fig. 5c**) (Venugopal et al., 2018). Secondly, we measured the 2^nd^ peak of the autocorrelation function of the membrane potential and show that the rescue of Na^+^ currents restored the rhythmicity to WT values in a dose-dependent manner (see **Fig. 5d** and **legend**). Lastly, Na^+^ current addition also reinstated the time intervals and duration to WT values (see **Fig. 5e** and **legend**).

**Figure 5.**
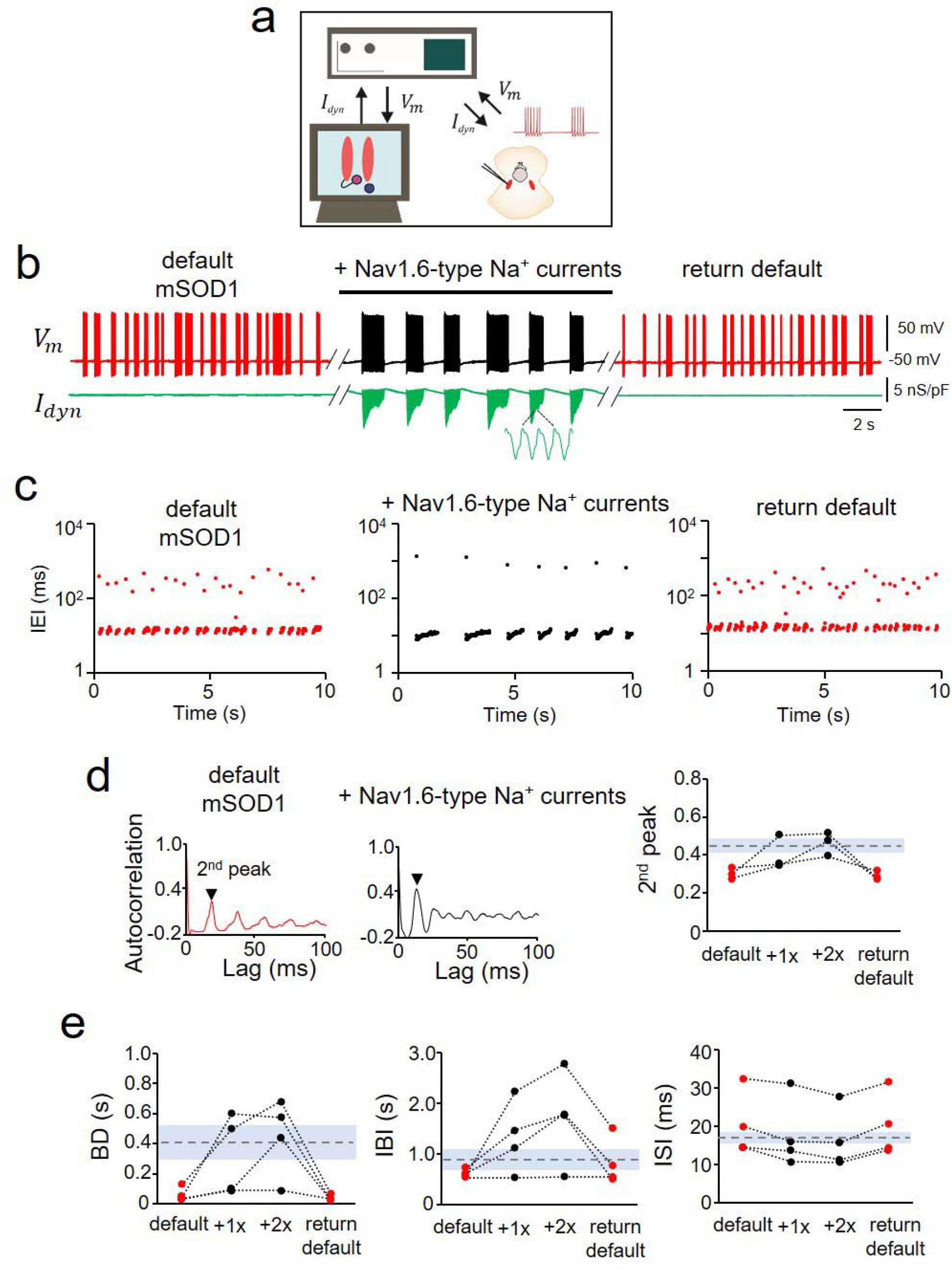
Rescue of Nav 1.6-type sodium currents in mSOD1 Mes V neurons using dynamic-clamp. **a.** Schematic shows the dynamic-clamp setup used to introduce conductance-based models of Nav1.6-type Na^+^ currents into mSOD1 Mes V neurons in real-time during whole-cell patch-clamp recording; *I_dyn_* is the computer-generated model Na^+^ current in combination with a step depolarization to drive the patched Mes V neuron; *V_m_* is the measured membrane voltage. **b.** Representative traces showing the membrane voltage in a bursting mSOD1 Mes V neuron with control/default behavior (*left red*), followed by addition of Nav1.6-type currents (*middle black*), that restores WT-like rhythmic bursting; subsequent removal of added currents returns default mSOD1 behavior (*right red*) in this neuron; Lower green trace shows *I_dyn_*. **c.** Time series plots of IEIs (log scale) for the three different conditions in (**b**); each dot represents an interval between two consecutive spikes (see detailed results). **d.** Autocorrelation function of the membrane voltage for default mSOD1 (*left*) and with addition of Nav1.6 currents (middle); the height of the 2^nd^ autocorrelation peak highlights rhythmicity (arrowhead); *Right:* measured 2^nd^ peak values for 4 mSOD1 cells are shown, with dashed horizontal line showing average WT values and shaded grey region indicates ±SD. **e.** Treatment effects on burst characteristics including burst duration (BD), inter-burst-intervals (IBI) and inter-spike-intervals within bursts (ISI) shown for various mSOD1 cells tested under the different conditions as in (**b**); dashed lines indicate average WT values with grey regions marking the ± s.d.

Taken together, the muscle spindle afferent neurons in the mSOD1 Mes V nucleus present early impairment in excitability including hypoexcitability and arrhythmicity which are associated with reductions in the voltage-gated Na^+^ currents and Nav1.6 Na^+^ channels.

### Mechanoreceptive and nociceptive trigeminal sensory neurons lack early excitability changes in the SOD1^G93A^ mouse

Next, to assess whether the altered excitability is restricted to the proprioceptive muscle spindle afferents in the mSOD1 Mes V nucleus, we examined the non-proprioceptive primary sensory neurons in the trigeminal ganglia (TG). The sensory architecture of the trigeminal system segregates the *Aα* -type proprioceptive neurons in the Mes V and the *Aβ* mechanoreceptors, *Aδ* nociceptors and C-fiber type neurons in the peripheral TG (Lazarov, 2002). Therefore, we examined whether the different types of sensory neurons in the TG were affected differently by SOD1 mutation. To conduct *in vitro* patch-clamp electrophysiology, we performed acute dissociation of the trigeminal ganglia in P8 – P14 WT and mSOD1 mice, age-matched with mice used to examine Mes V neurons (**Figs. 6a, b**) (also see **Methods Section Ib**). In both WT and mSOD1 TG neurons, we were able to distinguish the A*β* from the A*δ* nociceptive neurons based on action potential duration and presence of a hump as shown in **Fig. 6c** (Xu et al., 2010; Kim et al., 2011). A small proportion of C-fiber type neurons were also present in our dataset, which were distinguished from the A-type neurons based on a lack of membrane sag during the hyperpolarizing current injections (**Fig. 6c**: a white block arrow in the left two panels highlight membrane sag). However, these cells exhibited single spikes and were comparatively fewer in both WT and mSOD1 datasets to yield meaningful statistical assessment, and hence were excluded from further analysis. Interestingly, there were no proportional shifts in the sample dataset among these different types of neurons between WT and mSOD1 mice as shown in **Fig. 6d**. We compared the basic membrane properties and action potential characteristics of WT and mSOD1 TG neurons as shown in **Tables 3, 4**, the spike threshold current and voltage as shown in **Fig. 6e**. None of these properties were different between WT and mSOD1. The average spike frequency responses to increasing steps of depolarization also did not present any significant changes (see **Figs. 6f, g**). Lastly, the autocorrelation function of the membrane voltage (see **Fig. 6h**) did not show arrhythmicity in the mSOD1 TG neurons compared to WT. Note that the 2^nd^ peak of the autocorrelation function was similar and subsequent peaks were also similarly discernable with the autocorrelation time lag of 100 *ms*. These results suggest that non-proprioceptive sensory neurons are resistant to early excitability changes in the SOD1^G93A^ mouse model for ALS.

**TABLE 3.** Membrane properties of TG neurons.

|  | Aβ |  | Aδ |  | C |  |
| --- | --- | --- | --- | --- | --- | --- |
|  | WT (n=18) | mSOD1 (n=13) | WT (n=15) | mSOD1 (n=11) | WT (n=3) | mSOD1 (n=3) |
| RMP, mV | -53.2 ± 0.9 | -54.2 ± 0.9 | -50.8 ± 0.3 | -52.6 ± 1.1 | -62.5 ± 2.5 | -63.0 ± 3.0 |
| $R_{in}$, MΩ | 293.7 ± 86.4 | 164.6 ± 22.9 | 413.9 ± 50.1 | 373.2 ± 56.8 | 224.1 ± 50.6 | 308.8 ± 123.4 |
| $C_m$, pF | 32.8 ± 4.6 | 35.1 ± 4.2 | 20.3 ± 3.7 | 18.4 ± 4.8 | 21.8 ± 4.2 | 27.4 ± 8.8 |

**TABLE 4.**
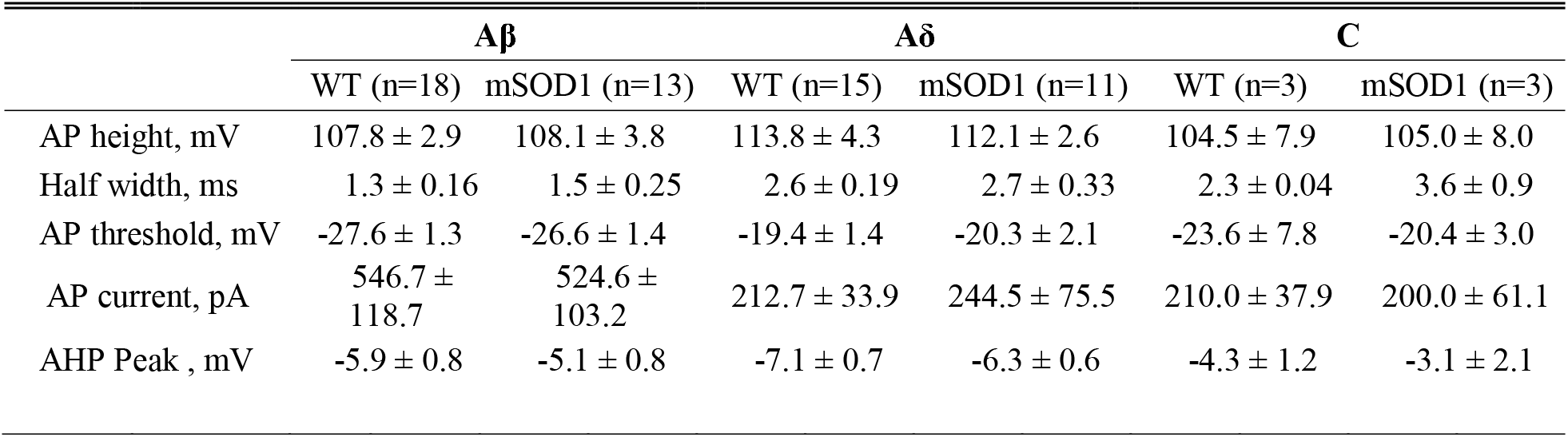
Action potential properties of TG neurons.

|  | Aβ |  | Aδ |  | C |  |
| --- | --- | --- | --- | --- | --- | --- |
|  | WT (n=18) | mSOD1 (n=13) | WT (n=15) | mSOD1 (n=11) | WT (n=3) | mSOD1 (n=3) |
| AP height, mV | 107.8 ± 2.9 | 108.1 ± 3.8 | 113.8 ± 4.3 | 112.1 ± 2.6 | 104.5 ± 7.9 | 105.0 ± 8.0 |
| Half width, ms | 1.3 ± 0.16 | 1.5 ± 0.25 | 2.6 ± 0.19 | 2.7 ± 0.33 | 2.3 ± 0.04 | 3.6 ± 0.9 |
| AP threshold, mV | -27.6 ± 1.3 | -26.6 ± 1.4 | -19.4 ± 1.4 | -20.3 ± 2.1 | -23.6 ± 7.8 | -20.4 ± 3.0 |
| AP current, pA | 546.7 ± 118.7 | 524.6 ± 103.2 | 212.7 ± 33.9 | 244.5 ± 75.5 | 210.0 ± 37.9 | 200.0 ± 61.1 |
| AHP Peak , mV | -5.9 ± 0.8 | -5.1 ± 0.8 | -7.1 ± 0.7 | -6.3 ± 0.6 | -4.3 ± 1.2 | -3.1 ± 2.1 |

**Figure 6.**
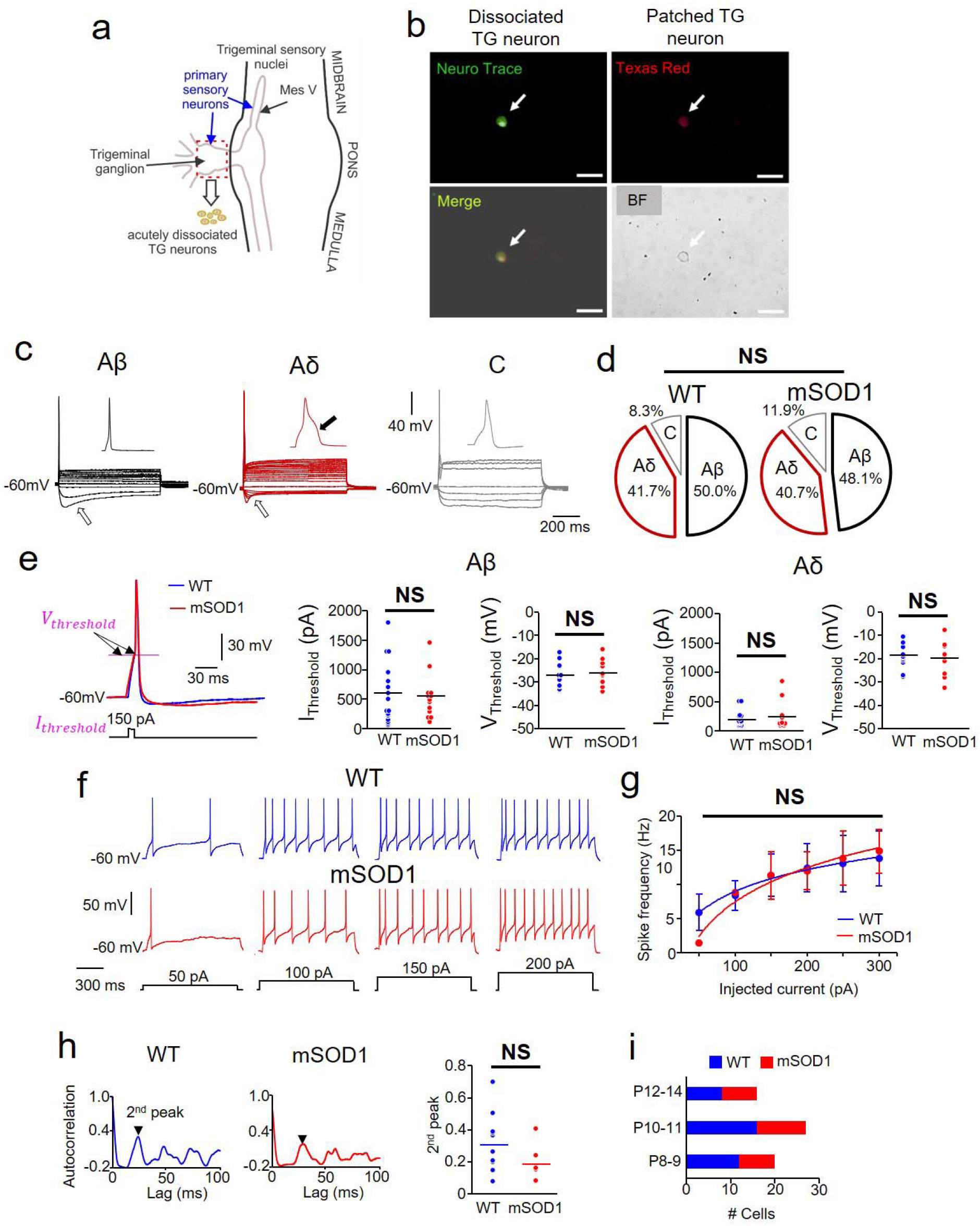
Excitability of primary sensory neurons in the TG of mSOD1 mice. **a.** Schematic showing the trigeminal sensory nuclei along the midbrain-brainstem regions; Primary 1a afferent neurons in the TG (red dashed rectangle) were acutely dissociated and whole-cell current-clamp recordings were performed. **b.** *Top left:* Example showing NeuN stain (green) used to identify the dissociated TG neurons; *Top right:* Example of a dissociated TG neuron filled with Texas red 568 dye during whole-cell recording; *Bottom left:* Merged image; *Bottom right:* Bright field (BF) image; Scale bars are 50 μm. **c.** TG neurons were classified into three types: Aβ, Aδ and C (see results for criteria); open arrows highlight membrane sag during a 1s hyperpolarizing step pulse in Aβ and Aδ types; black arrow highlights a hump in Aδ action potential. **d.** Percentage of the subtypes of TG neurons in WT and mSOD1 mice are not significantly different (NS). **e.** Inset shows the current (*I_Threshold_*) and voltage (*V_Threshold_*) thresholds for action potentials in WT (blue) and mSOD1 (red) TG neurons; Dot plots show comparison of these properties between WT (blue) and mSOD1 (red) within the Aβ and Aδ groups with no statistical significance (NS). **f.** Representative membrane voltage responses to increasing levels of 1s current injection (bottom traces) in repetitively firing WT and mSOD1 TG neurons (both Aδ-type); **g.** Spike frequency – Injected current responses in all the Aδ TG neurons that showed repetitive firing; No statistical significance (NS). **h.** Autocorrelation function of membrane potential in WT (*left panel*) and mSOD1 (*middle panel*) TG neurons; 2^nd^ peaks (arrows) are highlighted. *Right:* Dot plots showing the 2^nd^ peaks for all the cells with no significant difference (NS) between WT (blue) and mSOD1 (red) TG neurons. **i.** Bar charts summarize the age distribution of the mice including 8 WT mice (*n = 36*) and 7 mSOD1 mice (*n = 27*), where *n* is the number of cells for all the data presented in panels **e – h**.

**Figure 7.**
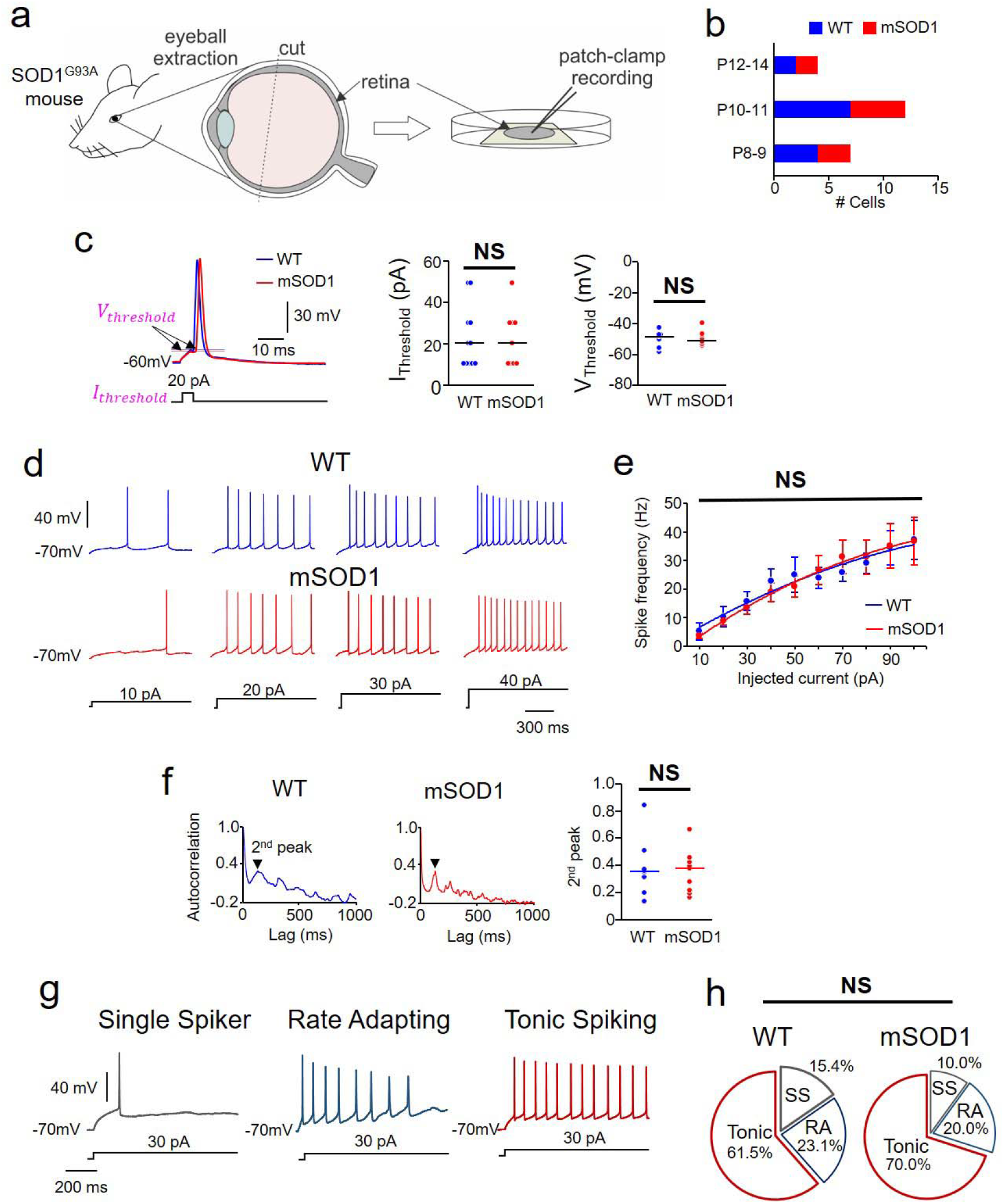
Excitability of primary sensory neurons in the retina of mSOD1 mice. **a.** Schematic showing retina extraction for patch-clamp recording. **b.** Bar charts summarize the age distribution of the mice including 7 WT mice (*n = 13*) and 5 mSOD1 mice (*n = 10*), where *n* is the number of cells for all the data presented in panels **c – h**. **c.** Inset shows the current (*I_Threshold_*) and voltage (*V_Threshold_*) thresholds for action potentials in WT (blue) and mSOD1 (red) RG neurons; Dot plots show comparison of these properties between WT (blue) and mSOD1 (red) RG neurons tested; NS indicates no statistical significance. **f.** Representative membrane voltage responses to increasing levels of 1s current injection (bottom traces) in repetitively firing WT and mSOD1 RG neurons; **g.** Spike frequency – Injected current responses in all the RG neurons that showed repetitive firing; No statistical significance (NS). **h.** Autocorrelation function of membrane potential in WT (*left panel*) and mSOD1 (*middle panel*) RG neurons; 2^nd^ peaks (arrows) are highlighted. *Right:* Dot plots showing the 2^nd^ peaks for all the cells with no significant difference (NS) between WT (blue) and mSOD1 (red) TG neurons. **g.** RG neurons were classified into three types: Single spikers (SS), Rate Adapting (RA) and Tonic Spiking (see results for details). **d.** Percentage of the subtypes of RG neurons in WT and mSOD1 mice are not significantly different (NS).

### Early excitability changes are absent in the SOD1^G93A^ retinal ganglion neurons

Lastly, we investigated whether the peripheral visual sensory retinal ganglion neurons (RGNs) show altered excitability in the mSOD1 mice. For these experiments, we used a whole retinal preparation and conducted whole-cell patch-clamp electrophysiology *in situ* in the RGNs in P8 – P14 mice (**Fig. 7a, b;** also see **Methods Section Ic**). The rationale for examining RGN excitability was three-fold: **1)** to test whether functional abnormalities were limited to muscle spindle afferent proprioceptive neurons while absent in the visual sensory neurons, similar to the mechanoreceptive and nociceptive TG neurons, **2)** to ascertain whether excitability changes were not confounded by the type of preparation (*in vitro* dissociated TGNs versus *in situ* RGNs), and, **3)** to clarify whether postnatal excitability changes in other sensory neurons are equally immune in parallel with maturing peripheral sensory systems in mice (Cabanes et al., 2002; Chen et al., 2009). Similar to the Mes V and TG neurons, we compared the membrane properties (**Tables 5, 6**), action potential thresholds (**Fig. 7c**), spike frequency – injected current responses (**Fig. 7d, e**), rhythmicity/regularity of membrane voltage spikes (**Fig. 7f**), and, proportional distribution of apparent electrophysiological subtypes of RGNs in the WT and mSOD1 mice (**Figs. 7g, h**). We found that none of the above properties tested were different between the two groups. These results confirm that the early excitability changes are limited to muscle spindle afferent proprioceptive neurons in the cranial sensory system of the SOD1^G93A^ mouse model for ALS.

**TABLE 5.** Membrane properties of RGCs.

|  | WT (n=13) | SOD1 (n=10) |
| --- | --- | --- |
| RMP, mV | -55.4 ± 2.0 | -54.1 ± 2.4 |
| $R_{in}$, MΩ | 462.3 ± 19.3 | 436.2 ± 24.4 |
| $C_m$, pF | 29.0 ± 3.3 | 29.3 ± 4.4 |

**TABLE 6.** Action potential properties of RGCs.

|  | WT (n=13) | mSOD1 (n=10) |
| --- | --- | --- |
| AP height, mV | 67.6 ± 3.9 | 66.3 ± 5.3 |
| Half width, ms | 3.8 ± 0.5 | 4.0 ± 0.7 |
| AP threshold, mV | -50.6 ± 1.5 | -51.6 ± 2.2 |
| AP current, pA | 22.7 ± 4.7 | 22.2 ± 4.6 |
| AHP Peak , mV | -6.1 ± 0.4 | -6.4 ± 0.8 |

### Computer modeling predicts motor dysfunction due to perturbed proprioceptive sensory gating

Difficulty in chewing and swallowing is a common clinical feature observed in ALS patients with both spinal and bulbar onset (Riera-Punet et al., 2018a; Riera-Punet et al., 2018b). In the SOD1^G93A^ mouse model, mastication is severely impaired early during disease development (Lever et al., 2009). During normal mastication, the Mes V neurons relay sensory feedback from the jaw muscle spindles to adjust the activity and force generated in the jaw-closer muscles. Nearly 80% of the glutamatergic projections from the Mes V neurons synapse on the jaw-closer trigeminal motor pools and provide muscle stretch reflex inputs to the TMNs (Yoshida et al., 2017). It is likely that the observed abnormalities in the mSOD1 Mes V excitability alter the jaw stretch reflex control. To examine how irregular discharge in the mSOD1 Mes V neurons might modify motor discharge, we utilized a computational model of a simplified sensorimotor network. Our model consisted of a realistic sensory Mes V neuron which provides strong monosynaptic excitation to a postsynaptic trigeminal motor neuron (TMN) (Trueblood et al., 1996). We used an oversimplified, yet realistic construct, and tested how irregularities in ongoing burst patterns in the sensory Mes V neuron, such as during rhythmic jaw movements, could exclusively modulate the discharge patterns in a TMN. In **Fig. 8a**, we first illustrate rhythmic burst discharge such as in a WT Mes V neuron reproduced by the model. Such a regular pattern was converted into irregular/arrhythmic discharge observed in the mSOD1 Mes V neurons by a 25-50% reduction in the Mes V persistent and resurgent Na^+^ conductances and addition of sub-threshold stochastic inputs to enhance burst irregularities (see **Fig. 8b**) (also see (Venugopal et al., 2018)). We further assumed that the TMN model displays dendritic Ca^2+^ currents which can mediate plateau potentials and membrane bistability, often observed in brainstem and spinal MNs (Hsiao et al., 1998; Lee and Heckman, 1998; Hsiao et al., 2005). The model Mes V directly excited the dendritic compartment of the TMN which in turn depolarized the electrically coupled TMN soma. We further assumed that such depolarization drives dendritic Ca^2+^-mediated plateau potentials as would occur during NMDA receptor activation (Hsiao et al., 2002; Manuel et al., 2012). Activation of Ca^2+^ plateau depolarized the TMN soma and enabled sensorimotor synchronization driving downstream motor discharge (see **Fig. 8c**). However, when the sensory patterns were irregular, such synchronization was perturbed (see **Fig. 8d**). In particular, the shorter burst intervals mimicking mSOD1 discharge patterns induced instances of asynchronous self-sustained discharge in the TMN (see dashed boxes in **Fig. 8d**). In our model, this was due to inadequate deactivation of the plateau causing dendritic Ca^2+^ currents in the motor neuron between sensory bursts. Although under *in vivo* conditions, other mechanisms such as synaptic inhibition and Ca^2+^-activated K+ currents might effectively regulate Ca^2+^ plateaus and jaw reflexes (e.g., (Inoue et al., 1994; Hultborn et al., 2003; Li and Bennett, 2007; Venugopal et al., 2012)), our results highlight that rhythmic sensory timing, or lack thereof, can modulate sensorimotor synchronization. Taken together, one putative consequence of proprioceptive sensory irregularities leads to an asynchronous sustained motor discharge and may partly contribute to looming dysfunctions such as muscle fasciculations which are common symptoms in ALS (Hirota et al., 2000; Vucic and Kiernan, 2006) (see summary in **Fig. 8e**).

**Figure 8.**
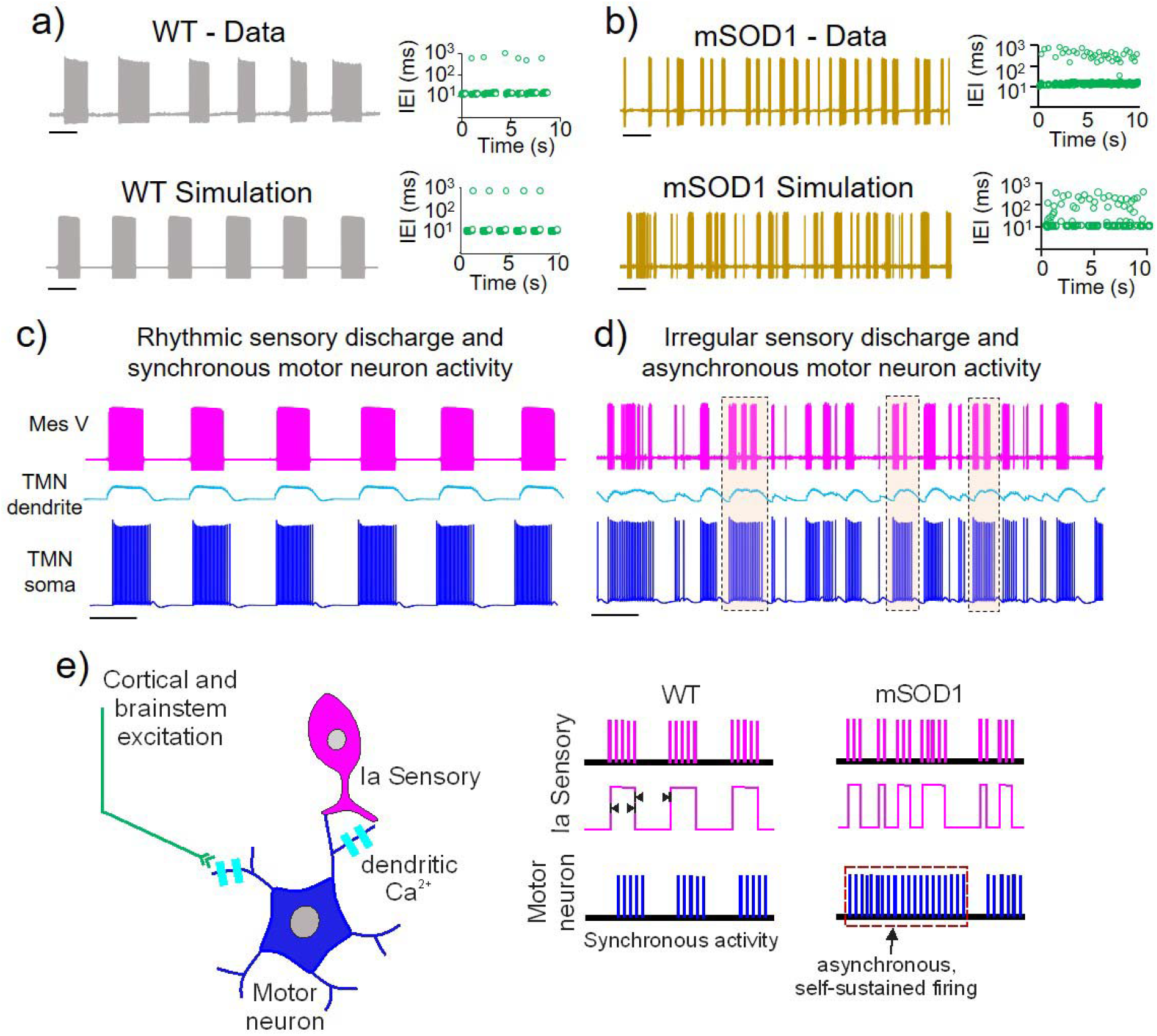
Computer-based model predicts irregular sensory driven motor asynchrony. **a)** Rhythmic burst pattern in a WT Mes V sensory neuron (top trace) is reproduced in a conductance-based Mes V neuron model (bottom); corresponding time series graphs of interevent intervals are shown in the top and bottom right panels (green circles). **b)** Burst irregularities in a mSOD1 Mes V neuron (top trace) is reproduced in the model neuron (bottom trace) by a 25% reduction in persistent Na^+^ conductance and a 50% reduction in resurgent Na^+^ conductance together with a slight increase in membrane potential noise; corresponding time series graphs of inter-event intervals are shown in the top and bottom right panels (green circles). **c, d)** Simulated post-synaptic membrane potentials in the motor dendrite (cyan) and soma (blue) for WT **(c)** and mSOD1 **(d)** sensory patterns, shown in the top magenta traces. Frequent occurrences of asynchronous self-sustained motor discharge patterns are highlighted by dashed rectangles. **e)** Schematic showing putative motor asynchrony and dysfunctional discharge patterns resulting from burst irregularities in the primary Ia afferent neuron.

### Discussion

In this study, we focused on SOD1^G93A^ mutation-induced modifications to proprioceptive sensory neurons (Mes V cells), at an early time point when MN dysregulation was previously reported (Venugopal et al., 2015). There are four main findings: 1) an early circuit-specific reduced excitability exclusively in the trigeminal proprioceptive sensory neurons in comparison with mechanoreceptive, nociceptive and visual sensory neurons in age-matched mSOD1 mice, 2) impaired bursting in the proprioceptive Mes V neurons associated with a down-regulation of Nav1.6 Na^+^ currents and ion channels, 3) rescue of normal burst patterns in the mSOD1 Mes V neurons upon restoration of Na^+^ currents, and, 4) computational model-based prediction of an effect of aberrant sensory gating on motor neuron discharge patterns. Considering these results, we discuss the consequences on disease development and progression.

### Circuit-specific vulnerability in ALS

Selective vulnerability is a hallmark of neurodegenerative diseases (Double et al., 2010); however, the factors which determine such selectivity are elusive and remain poorly understood. In ALS, as also with normal aging (Kanning et al., 2010), motor neurons (MNs) within a motor pool present selective and preferential vulnerability (Frey et al., 2000; Pun et al., 2006; Hegedus et al., 2007; Saxena and Caroni, 2011). Specifically, α-MNs forming the fast-fatigable motor units preferentially die followed by fast fatigue-resistant MNs, while the neighboring slow α-MNs and γ-MNs remain resistant to degeneration (Hegedus et al., 2007; Hegedus et al., 2008; Lalancette-Hebert et al., 2016). Although multiple intrinsic factors such as cell size (Dukkipati et al., 2018), Ca^2+^ buffering capacities (von Lewinski and Keller, 2005), synaptic organization (Nimchinsky et al., 2000; Lorenzo and Barbe, 2006), gene and protein expression patterns (Brockington et al., 2013; Comley et al., 2015) could govern selective vulnerability, the underlying network architecture can be a crucial determinant. For instance, a lack of muscle spindle afferent terminals on γ- and ocular MNs has been suggested as a mechanism of disease resistance (Keller and Robinson, 1971; Lalancette-Hebert et al., 2016). These afferents relay the proprioceptive sensory feedback which modulate motor neuron activity (the stretch reflex) during muscle force generation via glutamatergic excitation (e.g., (Chandler, 1989; Mentis et al., 2011)). Ablation of such spindle afferents significantly delayed MN death and disease progression in the SOD1^G93A^ mouse model (Lalancette-Hebert et al., 2016). Curiously, even in invertebrate SOD1-knock-in model systems, muscle spindle afferents act as an early trigger for MN degeneration (Held et al., 2019). This suggests the possibility that the proprioceptive feedback may indeed represent a phylogenetically conserved pathway of disease vulnerability. Furthermore, in a mouse model for Spinal Muscular Atrophy (SMA), a related motor neuron disease, proprioceptive inputs play a predominant role in triggering MN degeneration (Mentis et al., 2011). Advancing these findings, our comparative analysis of excitability of multiple sensory neurons revealed a SOD1 mutation-driven modification exclusively in the proprioceptive sensory neurons. Although non-motor and non-cell autonomous triggers have been implicated for MN death in ALS (Boillée et al., 2006a), to date, there is no direct evidence for exclusive circuit elements showing intrinsic electrophysiological abnormalities. In combination with our previous report on early dysregulation of MN excitability (Venugopal et al., 2015), the present results implicate a defective sensorimotor network in ALS, which may represent a convergent neuroanatomical pathway across multiple neurodegenerative motor neuron diseases.

### Excitability change – An early disease compensation and a marker for vulnerable circuits

Neurodegenerative diseases involve a protracted phase of progressive decline in the functional homeostasis of vulnerable neurons (Roselli and Caroni, 2015). In mouse models of ALS, hyperexcitability represents the earliest form of homeostatic disruption in brainstem, spinal and corticomotor neurons which are vulnerable to degeneration (Pieri et al., 2003; Amendola et al., 2007; Bories et al., 2007; van Zundert et al., 2008; Vucic et al., 2009; Quinlan et al., 2011; Venugopal et al., 2015). These results prompted a hyperexcitability-driven excitotoxicity hypothesis for cell death in ALS (Durand et al., 2006; van Zundert et al., 2012). Although attractive, a causal relationship between hyperexcitability and subsequent neurodegeneration is not well-supported (e.g., (Leroy et al., 2014; Simon et al., 2016)). Alternatively, it is likely that early hyperexcitability represents a compensatory mechanism of survival in normally low excitable cells since enhancing intrinsic excitability promotes neuroprotection (Saxena et al., 2013). Early compensation is also possible in slow MNs which normally show high excitability and are resistant to degeneration in ALS. Some of these slow MNs were hypoexcitable early in the SOD1^G93A^ mouse (Venugopal et al., 2015). Such *early* hypoexcitability in surviving MNs likely represents a way to moderate energy demands for spike generation which can be costly (e.g. (Le Masson et al., 2014)). This in turn could support cell survival in a disease background. A similar strategy could be used by the proprioceptive sensory neurons shown here which are involved in ongoing reflex control. We further speculate that hypoexcitable shift in the set point operation of neurons within a vulnerable circuitry could be triggered by disease-induced changes in astroglial cells which monitor extracellular ionic homeostasis (e.g. Ca^2+^ and K^+^) as well as bursting in Mes V neurons (Kadala et al., 2015; Morquette et al., 2015). Such crosstalk leading to compensatory intrinsic excitability changes needs to be further clarified. Notwithstanding a compensatory function, early excitability changes seem to be clear markers of vulnerable circuitry. For instance, disease resistant MNs (e.g., oculomotor neurons) (Venugopal et al., 2015) or non-proprioceptive sensory neurons as shown here do not present early modifications in excitability. In contrast a specific set of bursting proprioceptive Mes V neurons were hypoexcitable which form a monosynaptic jaw stretch reflex circuitry with vulnerable TMNs. Such reduced presynaptic sensory excitability could also be a trigger for compensatory hyperexcitability in the postsynaptic TMNs (Venugopal et al., 2015). However, it is yet unclear whether and how such sensory alterations contribute to MN degeneration (e.g., see (Lalancette-Hebert et al., 2016; Held et al., 2019)). Taken together, early excitability changes are crucial markers of vulnerable circuitry and may represent an early compensation to meet energy demands and likely support cell survival early during disease development.

### Altered activity patterns and ionic mechanisms as pathophysiological substrates

Altered phasic/burst patterns, reduced excitability and loss of excitation are defining features of vulnerability to neurodegeneration (Roselli and Caroni, 2015). Here we present a unique result highlighting early irregular and impaired burst discharge in proprioceptive sensory neurons forming a vulnerable sensorimotor circuitry. One consequence of irregular bursting could affect sensory gating of postsynaptic motor discharge as shown by our computational model. Secondly, impaired burst duration and frequencies together with reductions in persistent Na^+^ current could reduce extracellular Ca^2+^ and affect astrocyte-mediated Ca^2+^ buffering in the Mes V sensory nucleus (Morquette et al., 2015). Alternatively, reduced persistent Na^+^ and bursting could be a direct consequence of mSOD1 astrocyte dysfunction (Su et al., 2001). Curiously RGCs which also rely on persistent Na^+^ current for repetitive spiking (Boiko et al., 2003; Van Wart and Matthews, 2006a, b), did not show altered rhythmicity as measured by the autocorrelation functions. These observations suggest that the mutation could induce observed changes via altered extrinsic factors such as astrocyte dysfunction or by altered intrinsic factors such as abnormal protein synthesis contributing to the observed Na^+^ channelopathy in Mes V neurons. Furthermore, in the trigeminal system, the nonproprioceptive sensory neurons in the trigeminal ganglia were resistant to abnormalities in the SOD1^G93A^ mouse. Given that the trigeminal ganglia and Mes V neurons share ontogeny (Lazarov, 2002), an exclusive change only in Mes V supports extrinsic triggers within the jaw stretch reflex circuitry causing abnormal excitability.

Secondary effects of altered burst patterns could lead to recruitment of distinct types of voltage-dependent Ca^2+^ currents; resulting alterations in intracellular Ca^2+^ levels and kinetics can activate distinct CREB-dependent gene expression (Wheeler et al., 2012) and differentially activate nuclear translocation transcription factors such as the NFAT family, important in immune response (Hernández-Ochoa et al., 2007). Importantly, reduced excitability can deplete intracellular ER Ca^2+^ stores and disrupt protein homeostasis (e.g., a ubiquitin-dependent degradation of transcription factors important for synaptic plasticity (Lalonde et al., 2014)). Taken together, we suggest that identifying presymptomatic excitability changes, their upstream physiological triggers (e.g., astrocyte dysfunction) and downstream effectors (e.g., CREB-mediated gene expression changes) can elucidate the cascade of events leading to neurodegeneration. Additionally, excitability changes also represent early markers of disease vulnerability in ALS.

## Author Contributions

Study design and supervision (SV, SHC). SS conducted most of the experiments with TY and IS collaboration on trigeminal ganglia recordings, AP and RO collaboration on dynamic-clamp experiments. KQ conducted immunofluorescence experiments, MWP collaborated. SV conducted computational modeling and SHC collaborated. SS and SV analyzed the data and prepared the figures. SV wrote the manuscript; all the authors contributed to the final version.

## Acknowledgements

We are very grateful to Drs. Michael Levine, Sampath Alapakkam, Xia Yang and their lab members for their generous support on various aspects of this project. We thank Dr. Yatendra Mulpuri and Ravindu Udugampola for help with the histology experiments.

## References

Amendola J, Gueritaud JP, d’Incamps BL, Bories C, Liabeuf S, Allene C, Pambo-Pambo A, Durand J (2007) Postnatal electrical and morphological abnormalities in lumbar motoneurons from transgenic mouse models of amyotrophic lateral sclerosis. Arch Ital Biol 145:311–323.

Boiko T, Van Wart A, Caldwell JH, Levinson SR, Trimmer JS, Matthews G (2003) Functional specialization of the axon initial segment by isoform-specific sodium channel targeting. J Neurosci 23:2306–2313.

Boillée S, Vande Velde C, Cleveland DW (2006a) ALS: a disease of motor neurons and their nonneuronal neighbors. Neuron 52:39–59.

Boillée S, Yamanaka K, Lobsiger CS, Copeland NG, Jenkins NA, Kassiotis G, Kollias G, Cleveland DW (2006b) Onset and progression in inherited ALS determined by motor neurons and microglia. Science 312:1389–1392.

Booth V, Rinzel J, Kiehn O (1997) Compartmental Model of Vertebrate Motoneurons for Ca2+-Dependent Spiking and Plateau Potentials Under Pharmacological Treatment. Journal of Neurophysiology 78:3371–3385.

Bories C, Amendola J, Lamotte d’Incamps B, Durand J (2007) Early electrophysiological abnormalities in lumbar motoneurons in a transgenic mouse model of amyotrophic lateral sclerosis. Eur J Neurosci 25:451–459.

Brocard F, Verdier D, Arsenault I, Lund JP, Kolta A (2006) Emergence of intrinsic bursting in trigeminal sensory neurons parallels the acquisition of mastication in weanling rats. J Neurophysiol 96:2410–2424.

Brockington A, Ning K, Heath P, Wood E, Kirby J, Fusi N, Lawrence N, Wharton S, Ince P, Shaw P (2013) Unravelling the enigma of selective vulnerability in neurodegeneration: motor neurons resistant to degeneration in ALS show distinct gene expression characteristics and decreased susceptibility to excitotoxicity Acta Neuropathol 125:95–109.

Brownstone RM, Lancelin C (2018) Escape from homeostasis: spinal microcircuits and progression of amyotrophic lateral sclerosis. J Neurophysiol 119:1782–1794.

Bruijn LI, Miller TM, Cleveland DW (2004) Unraveling the mechanisms involved in motor neuron degeneration in ALS. Annu Rev Neurosci 27:723–749.

Cabanes C, López de Armentia M, Viana F, Belmonte C (2002) Postnatal changes in membrane properties of mice trigeminal ganglion neurons. J Neurophysiol 87:2398–2407.

Carlin K, Jones K, Jiang Z, Jordan L, Brownstone R (2000) Dendritic L-type calcium currents in mouse spinal motoneurons: implications for bistability. Eur J Neurosci 12:1635–1646.

Casas C, Manzano R, Vaz R, Osta R, Brites D (2016) Synaptic Failure: Focus in an Integrative View of ALS. Brain Plast 1:159–175.

Chandler SH (1989) Evidence for excitatory amino acid transmission between mesencephalic nucleus of V afferents and jaw-closer motoneurons in the guinea pig. Brain Res 477:252–264.

Chandler SH, Baker LL, Goldberg LJ (1984) Characterization of synaptic potentials in hindlimb extensor motoneurons during L-DOPA-induced fictive locomotion in acute and chronic spinal cats. Brain Res 303:91–100.

Chen L, Liu C, Liu L (2008) The modulation of voltage-gated potassium channels by anisotonicity in trigeminal ganglion neurons. Neuroscience 154:482–495.

Chen M, Weng S, Deng Q, Xu Z, He S (2009) Physiological properties of direction-selective ganglion cells in early postnatal and adult mouse retina. J Physiol 587:819–828.

Cleveland DW, Rothstein JD (2001) From Charcot to Lou Gehrig: deciphering selective motor neuron death in ALS. Nat Rev Neurosci 2:806–819.

Comley L, Allodi I, Nichterwitz S, Nizzardo M, Simone C, Corti S, Hedlund E (2015) Motor neurons with differential vulnerability to degeneration show distinct protein signatures in health and ALS. Neuroscience 291:216–229.

Del Negro CA, Chandler SH (1997) Physiological and theoretical analysis of K+ currents controlling discharge in neonatal rat mesencephalic trigeminal neurons. J Neurophysiol 77:537–553.

Del Negro CA, Chandler SH (1998) Regulation of intrinsic and synaptic properties of neonatal rat trigeminal motoneurons by metabotropic glutamate receptors. J Neurosci 18:9216–9226.

Do MT, Bean BP (2003) Subthreshold sodium currents and pacemaking of subthalamic neurons: modulation by slow inactivation. Neuron 39:109–120.

Double KL, Reyes S, Werry EL, Halliday GM (2010) Selective cell death in neurodegeneration: why are some neurons spared in vulnerable regions? Prog Neurobiol 92:316–329.

Dukkipati SS, Garrett TL, Elbasiouny SM (2018) The vulnerability of spinal motoneurons and soma size plasticity in a mouse model of amyotrophic lateral sclerosis. J Physiol 596:1723–1745.

Durand J, Amendola J, Bories C, Lamotte d’Incamps B (2006) Early abnormalities in transgenic mouse models of amyotrophic lateral sclerosis. J Physiol Paris 99:211–220.

Enomoto A, Han JM, Hsiao CF, Chandler SH (2007) Sodium currents in mesencephalic trigeminal neurons from Nav1.6 null mice. J Neurophysiol 98:710–719.

Enomoto A, Han JM, Hsiao CF, Wu N, Chandler SH (2006) Participation of sodium currents in burst generation and control of membrane excitability in mesencephalic trigeminal neurons. J Neurosci 26:3412–3422.

Ermentrout B (2001) XPPAUT5.0 – the differential equations tool. In.

Ferrucci M, Spalloni A, Bartalucci A, Cantafora E, Fulceri F, Nutini M, Longone P, Paparelli A, Fornai F (2010) A systematic study of brainstem motor nuclei in a mouse model of ALS, the effects of lithium. Neurobiol Dis 37:370–383.

Frey D, Schneider C, Xu L, Borg J, Spooren W, Caroni P (2000) Early and selective loss of neuromuscular synapse subtypes with low sprouting competence in motoneuron diseases. J Neurosci 20:2534–2542.

Gupta A, Elgammal FS, Proddutur A, Shah S, Santhakumar V (2012) Decrease in tonic inhibition contributes to increase in dentate semilunar granule cell excitability after brain injury. J Neurosci 32:2523–2537.

Heckman C, Gorassini M, Bennett D (2005) Persistent inward currents in motoneuron dendrites: implications for motor output. Muscle Nerve 31:153–156.

Heckman CJ, Binder MD (1991) Analysis of Ia-inhibitory synaptic input to cat spinal motoneurons evoked by vibration of antagonist muscles. J Neurophysiol 66:1888–1893.

Hedlund E, Karlsson M, Osborn T, Ludwig W, Isacson O (2010) Global gene expression profiling of somatic motor neuron populations with different vulnerability identify molecules and pathways of degeneration and protection. Brain 133:2313–2330.

Hegedus J, Putman CT, Gordon T (2007) Time course of preferential motor unit loss in the SOD1 G93A mouse model of amyotrophic lateral sclerosis. Neurobiol Dis 28:154–164.

Hegedus J, Putman CT, Tyreman N, Gordon T (2008) Preferential motor unit loss in the SOD1 G93A transgenic mouse model of amyotrophic lateral sclerosis. J Physiol 586:3337–3351.

Held A, Major P, Sahin A, Reenan RA, Lipscombe D, Wharton KA (2019) Circuit Dysfunction in. J Neurosci 39:2347–2364.

Henderson G, Pepper CM, Shefner SA (1982) Electrophysiological properties of neurons contained in the locus coeruleus and mesencephalic nucleus of the trigeminal nerve in vitro. Exp Brain Res 45:29–37.

Hernández-Ochoa EO, Contreras M, Cseresnyés Z, Schneider MF (2007) Ca2+ signal summation and NFATc1 nuclear translocation in sympathetic ganglion neurons during repetitive action potentials. Cell Calcium 41:559–571.

Hirota N, Eisen A, Weber M (2000) Complex fasciculations and their origin in amyotrophic lateral sclerosis and Kennedy’s disease. Muscle Nerve 23:1872–1875.

Hodgkin AL, Huxley AF (1952) A quantitative description of membrane current and its application to conduction and excitation in nerve. J Physiol 117:500–544.

Hsiao CF, Wu N, Chandler SH (2005) Voltage-dependent calcium currents in trigeminal motoneurons of early postnatal rats: modulation by 5-HT receptors. J Neurophysiol 94:2063–2072.

Hsiao CF, Del Negro CA, Trueblood PR, Chandler SH (1998) Ionic basis for serotonin-induced bistable membrane properties in guinea pig trigeminal motoneurons. J Neurophysiol 79:2847–2856.

Hsiao CF, Wu N, Levine MS, Chandler SH (2002) Development and serotonergic modulation of NMDA bursting in rat trigeminal motoneurons. J Neurophysiol 87:1318–1328.

Hultborn H, Denton M, Wienecke J, Nielson J (2003) Variable amplification of synaptic input to cat spinal motoneurons by persistent inward current. Journal of Physiology 552:945–952.

Inoue T, Chandler SH, Goldberg LJ (1994) Neuropharmacological mechanisms underlying rhythmical discharge in trigeminal interneurons during fictive mastication. J Neurophysiol 71:2061–2073.

Iwata M, Hirano A (1978) Sparing of the Onufrowicz nucleus in sacral anterior horn lesions. Ann Neurol 4:245–249.

Kadala A, Verdier D, Morquette P, Kolta A (2015) Ion Homeostasis in Rhythmogenesis: The Interplay Between Neurons and Astroglia. Physiology (Bethesda) 30:371–388.

Kanning KC, Kaplan A, Henderson CE (2010) Motor neuron diversity in development and disease. Annu Rev Neurosci 33:409–440.

Keller EL, Robinson DA (1971) Absence of a stretch reflex in extraocular muscles of the monkey. J Neurophysiol 34:908–919.

Kim HY, Chung G, Jo HJ, Kim YS, Bae YC, Jung SJ, Kim JS, Oh SB (2011) Characterization of dental nociceptive neurons. J Dent Res 90:771–776.

Lalancette-Hebert M, Sharma A, Lyashchenko AK, Shneider NA (2016) Gamma motor neurons survive and exacerbate alpha motor neuron degeneration in ALS. Proc Natl Acad Sci U S A 113:E8316–E8325.

Lalonde J, Saia G, Gill G (2014) Store-operated calcium entry promotes the degradation of the transcription factor Sp4 in resting neurons. Sci Signal 7:ra51.

Lazarov NE (2002) Comparative analysis of the chemical neuroanatomy of the mammalian trigeminal ganglion and mesencephalic trigeminal nucleus. Prog Neurobiol 66:19–59.

Le Masson G, Przedborski S, Abbott L (2014) A Computational Model of Motor Neuron Degeneration. Neuron 83:975–988.

Lee R, Heckman C (1998) Bistability in spinal motoneurons in vivo: systematic variations in persistent inward currents. Journal of Neurophysiology 80:583–593.

Lee RH, Kuo JJ, Jiang MC, Heckman CJ (2003) Influence of active dendritic currents on input-output processing in spinal motoneurons in vivo. J Neurophysiol 89:27–39.

Lee Y, Morrison BM, Li Y, Lengacher S, Farah MH, Hoffman PN, Liu Y, Tsingalia A, Jin L, Zhang PW, Pellerin L, Magistretti PJ, Rothstein JD (2012) Oligodendroglia metabolically support axons and contribute to neurodegeneration. Nature 487:443–448.

Leroy F, Lamotte d’Incamps B, Imhoff-Manuel RD, Zytnicki D (2014) Early intrinsic hyperexcitability does not contribute to motoneuron degeneration in amyotrophic lateral sclerosis. Elife 3.

Lever TE, Gorsek A, Cox KT, O’Brien KF, Capra NF, Hough MS, Murashov AK (2009) An animal model of oral dysphagia in amyotrophic lateral sclerosis. Dysphagia 24:180–195.

Li X, Bennett DJ (2007) Apamin-sensitive calcium-activated potassium currents (SK) are activated by persistent calcium currents in rat motoneurons. J Neurophysiol 97:3314–3330.

Lin RJ, Bettencourt J, Wha Ite J, Christini DJ, Butera RJ (2010) Real-time experiment interface for biological control applications. Conf Proc IEEE Eng Med Biol Soc 2010:4160–4163.

Lorenzo L, Barbe A (2006) Differential expression of GABAA and glycine receptors in ALS-resistant vs. ALS-vulnerable motoneurons: possible implications for selective vulnerability of motoneurons. European Journal of Neuroscience 23:3161–3170.

Malin SA, Davis BM, Molliver DC (2007) Production of dissociated sensory neuron cultures and considerations for their use in studying neuronal function and plasticity. Nat Protoc 2:152–160.

Manuel M, Li Y, Elbasiouny SM, Murray K, Griener A, Heckman CJ, Bennett DJ (2012) NMDA induces persistent inward and outward currents that cause rhythmic bursting in adult rodent motoneurons. J Neurophysiol 108:2991–2998.

Marchenkova A, van den Maagdenberg AM, Nistri A (2016) Loss of inhibition by brain natriuretic peptide over P2X3 receptors contributes to enhanced spike firing of trigeminal ganglion neurons in a mouse model of familial hemiplegic migraine type-1. Neuroscience 331:197–205.

Margolis DJ, Detwiler PB (2007) Different mechanisms generate maintained activity in ON and OFF retinal ganglion cells. J Neurosci 27:5994–6005.

Mentis GZ, Blivis D, Liu W, Drobac E, Crowder ME, Kong L, Alvarez FJ, Sumner CJ, O’Donovan MJ (2011) Early functional impairment of sensory-motor connectivity in a mouse model of spinal muscular atrophy. Neuron 69:453–467.

Morquette P, Verdier D, Kadala A, Féthière J, Philippe AG, Robitaille R, Kolta A (2015) An astrocyte-dependent mechanism for neuronal rhythmogenesis. Nat Neurosci 18:844–854.

Murphy GJ, Rieke F (2006) Network variability limits stimulus-evoked spike timing precision in retinal ganglion cells. Neuron 52:511–524.

Nijssen J, Comley LH, Hedlund E (2017) Motor neuron vulnerability and resistance in amyotrophic lateral sclerosis. Acta Neuropathol 133:863–885.

Nimchinsky EA, Young WG, Yeung G, Shah RA, Gordon JW, Bloom FE, Morrison JH, Hof PR (2000) Differential vulnerability of oculomotor, facial, and hypoglossal nuclei in G86R superoxide dismutase transgenic mice. J Comp Neurol 416:112–125.

Pieri M, Albo F, Gaetti C, Spalloni A, Bengtson CP, Longone P, Cavalcanti S, Zona C (2003) Altered excitability of motor neurons in a transgenic mouse model of familial amyotrophic lateral sclerosis. Neurosci Lett 351:153–156.

Powers RK, Elbasiouny SM, Rymer WZ, Heckman CJ Contribution of Intrinsic Properties and Synaptic Inputs to Motoneuron Discharge Patterns: A Simulation Study. J Neurophysiol.

Puls I, Jonnakuty C, LaMonte BH, Holzbaur EL, Tokito M, Mann E, Floeter MK, Bidus K, Drayna D, Oh SJ, Brown RH, Ludlow CL, Fischbeck KH (2003) Mutant dynactin in motor neuron disease. Nat Genet 33:455–456.

Pun S, Santos AF, Saxena S, Xu L, Caroni P (2006) Selective vulnerability and pruning of phasic motoneuron axons in motoneuron disease alleviated by CNTF. Nat Neurosci 9:408–419.

Qu J, Myhr KL (2011) The morphology and intrinsic excitability of developing mouse retinal ganglion cells. PLoS One 6:e21777.

Quinlan KA, Schuster JE, Fu R, Siddique T, Heckman CJ (2011) Altered postnatal maturation of electrical properties in spinal motoneurons in a mouse model of amyotrophic lateral sclerosis. J Physiol 589:2245–2260.

Raman IM, Bean BP (2001) Inactivation and recovery of sodium currents in cerebellar Purkinje neurons: evidence for two mechanisms. Biophys J 80:729–737.

Rieke F, Warland D, van Steveninck RdR, Bialek W (1999) Spikes: Exploring the Neural Code, First Edition. Cambridge, Massachusetts, USA: MIT Press.

Riera-Punet N, Martinez-Gomis J, Paipa A, Povedano M, Peraire M (2018a) Alterations in the Masticatory System in Patients with Amyotrophic Lateral Sclerosis. J Oral Facial Pain Headache 32:84–90.

Riera-Punet N, Martinez-Gomis J, Willaert E, Povedano M, Peraire M (2018b) Functional limitation of the masticatory system in patients with bulbar involvement in amyotrophic lateral sclerosis. J Oral Rehabil 45:204–210.

Roselli F, Caroni P (2015) From intrinsic firing properties to selective neuronal vulnerability in neurodegenerative diseases. Neuron 85:901–910.

Saxena S, Caroni P (2011) Selective neuronal vulnerability in neurodegenerative diseases: from stressor thresholds to degeneration. Neuron 71:35–48.

Saxena S, Roselli F, Singh K, Leptien K, Julien J-P, Gros-Louis F, Caroni P (2013) Neuroprotection through excitability and mTOR required in ALS motoneurons to delay disease and extend survival. Neuron 80:80–96.

Schindelin J, Arganda-Carreras I, Frise E, Kaynig V, Longair M, Pietzsch T, Preibisch S, Reuden C, Saalfeld S, Schmid B, Tinevez J, White D, Hartenstein V, Eliceiri K, Tomancak P, Cardona A (2012) Fiji: an open-source platform for biological image analysis. Nature Methods 9:676–682.

Schneider C, Rasband W, Eliceiri K (2012) NIH Image to ImageJ: 25 years of image analysis. Nature Methods 9:671–675.

Schurr A, West CA, Rigor BM (1988) Lactate-supported synaptic function in the rat hippocampal slice preparation. Science 240:1326–1328.

Seki S, Chandler H S, Maheshwary R, Nisly H, Sampath P A, Olcese R, Wiedau-Pazos M, Venugopal S (2017) Pre-symptomatic abnormalities and associated channelopathies in spindle afferent trigeminal mesencephalic V neurons in a SOD1G93A mouse model for Amyotrophic Lateral Sclerosis. In: Society for Neuroscience 2017. Washington D.C.

Simon CM, Janas AM, Lotti F, Tapia JC, Pellizzoni L, Mentis GZ (2016) A Stem Cell Model of the Motor Circuit Uncouples Motor Neuron Death from Hyperexcitability Induced by SMN Deficiency. Cell Rep 16:1416–1430.

Spencer RF, Porter JD (1988) Structural organization of the extraocular muscles. Rev Oculomot Res 2:33–79.

Su H, Alroy G, Kirson ED, Yaari Y (2001) Extracellular calcium modulates persistent sodium current-dependent burst-firing in hippocampal pyramidal neurons. J Neurosci 21:4173–4182.

Taylor JP, Brown RH, Cleveland DW (2016) Decoding ALS: from genes to mechanism. Nature 539:197–206.

Theiss RD, Kuo JJ, Heckman CJ (2007) Persistent inward currents in rat ventral horn neurones. J Physiol 580:507–522.

Trueblood PR, Levine MS, Chandler SH (1996) Dual-component excitatory amino acid-mediated responses in trigeminal motoneurons and their modulation by serotonin in vitro. J Neurophysiol 76:2461–2473.

Turman JE (2007) The development of mastication in rodents: from neurons to behaviors. Arch Oral Biol 52:313–316.

Turman JE, Jr., Ajdari J, Chandler SH (1999) NMDA receptor NR1 and NR2A/B subunit expression in trigeminal neurons during early postnatal development. J Comp Neurol 409:237–249.

Turman JE, Jr., MacDonald AS, Pawl KE, Bringas P, Chandler SH (2000) AMPA receptor subunit expression in trigeminal neurons during postnatal development. J Comp Neurol 427:109–123.

Van Wart A, Matthews G (2006a) Expression of sodium channels Nav1.2 and Nav1.6 during postnatal development of the retina. Neurosci Lett 403:315–317.

Van Wart A, Matthews G (2006b) Impaired firing and cell-specific compensation in neurons lacking nav1.6 sodium channels. J Neurosci 26:7172–7180.

van Zundert B, Izaurieta P, Fritz E, Alvarez FJ (2012) Early pathogenesis in the adult-onset neurodegenerative disease amyotrophic lateral sclerosis. J Cell Biochem 113:3301–3312.

van Zundert B, Peuscher MH, Hynynen M, Chen A, Neve RL, Brown RH, Jr., Constantine-Paton M, Bellingham MC (2008) Neonatal neuronal circuitry shows hyperexcitable disturbance in a mouse model of the adult-onset neurodegenerative disease amyotrophic lateral sclerosis. J Neurosci 28:10864–10874.

Venugopal S, Hamm T, Jung R (2012) Differential contributions of somatic and dendritic calcium-dependent potassium currents to the control of motoneuron excitability following spinal cord injury Cognitive Neurodynamics 6:283–293.

Venugopal S, Hamm TM, Crook SM, Jung R (2011a) Modulation of inhibitory strength and kinetics facilitates regulation of persistent inward currents and motoneuron excitability following spinal cord injury. J Neurophysiol 106:2167–2179.

Venugopal S, Crook S, S S, R J (2011b) Principles of Computational Neuroscience. In: Biohybrid Systems – Nerves, Interfaces and Machines (R J, ed), pp 1 – 12: Wiley & Sons.

Venugopal S, Hsiao C, Sonoda T, Weidau-Pazos M, Chandler S (2015) Homeostatic dysregulation in membrane properties of masticatory motoneurons compared to oculomotor neurons in a mouse model for Amyotrophic Lateral Sclerosis. Journal of Neuroscience 35:707–720.

Venugopal S, Seki S, Terman D, Pantazis A, Olcese R, Wiedau-Pazos M, Chandler S (2018) A Mechanism of Real-Time Noise Modulation in Neurons. In: bioRXiv: Cold Spring Harbor Laboratory.

von Lewinski F, Keller BU (2005) Ca2+, mitochondria and selective motoneuron vulnerability: implications for ALS. Trends Neurosci 28:494–500.

Vucic S, Kiernan MC (2006) Axonal excitability properties in amyotrophic lateral sclerosis. Clin Neurophysiol 117:1458–1466.

Vucic S, Cheah BC, Kiernan MC (2009) Defining the mechanisms that underlie cortical hyperexcitability in amyotrophic lateral sclerosis. Exp Neurol 220:177–182.

Wang GY, Ratto G, Bisti S, Chalupa LM (1997) Functional development of intrinsic properties in ganglion cells of the mammalian retina. J Neurophysiol 78:2895–2903.

Wheeler DG, Groth RD, Ma H, Barrett CF, Owen SF, Safa P, Tsien RW (2012) Ca(V)1 and Ca(V)2 channels engage distinct modes of Ca(2+) signaling to control CREB-dependent gene expression. Cell 149:1112–1124.

Williamson TL, Cleveland DW (1999) Slowing of axonal transport is a very early event in the toxicity of ALS-linked SOD1 mutants to motor neurons. Nat Neurosci 2:50–56.

Wu N, Hsiao CF, Chandler SH (2001) Membrane resonance and subthreshold membrane oscillations in mesencephalic V neurons: participants in burst generation. J Neurosci 21:3729–3739.

Wu N, Enomoto A, Tanaka S, Hsiao CF, Nykamp DQ, Izhikevich E, Chandler SH (2005) Persistent sodium currents in mesencephalic v neurons participate in burst generation and control of membrane excitability. J Neurophysiol 93:2710–2722.

Xu S, Ono K, Inenaga K (2010) Electrophysiological and chemical properties in subclassified acutely dissociated cells of rat trigeminal ganglion by current signatures. J Neurophysiol 104:3451–3461.

Yamamoto T, Ono K, Hitomi S, Harano N, Sago T, Yoshida M, Nunomaki M, Shiiba S, Watanabe S, Nakanishi O, Inenaga K (2013) Endothelin Receptor-mediated Responses in Trigeminal Ganglion Neurons Journal of Dental Research: 1–5.

Yamanaka K, Chun SJ, Boillee S, Fujimori-Tonou N, Yamashita H, Gutmann DH, Takahashi R, Misawa H, Cleveland DW (2008) Astrocytes as determinants of disease progression in inherited amyotrophic lateral sclerosis. Nat Neurosci 11:251–253.

Yang J, Xing JL, Wu NP, Liu YH, Zhang CZ, Kuang F, Han VZ, Hu SJ (2009) Membrane current-based mechanisms for excitability transitions in neurons of the rat mesencephalic trigeminal nuclei. Neuroscience 163:799–810.

Yoshida A, Moritani M, Nagase Y, Bae YC (2017) Projection and synaptic connectivity of trigeminal mesencephalic nucleus neurons controlling jaw reflexes. J Oral Sci 59:177–182.

